# Lrig1 and Wnt dependent niches dictate segregation of resident immune cells and melanocytes in murine tail epidermis

**DOI:** 10.1101/2021.12.22.473845

**Authors:** Susanne C. Baess, Ann-Kathrin Burkhart, Sabrina Cappello, Annika Graband, Kristin Seré, Martin Zenke, Catherin Niemann, Sandra Iden

## Abstract

The barrier-forming, self-renewing mammalian epidermis comprises keratinocytes, pigment-producing melanocytes, and resident immune cells as first-line host defense. In murine tail skin, interfollicular epidermis patterns into pigmented ′scale′ and hypopigmented ′interscale′ epidermis. Why and how mature melanocytes accumulate in scale epidermis is unresolved. Here, we delineate a cellular hierarchy among epidermal cell types that determines skin patterning. Already during postnatal development, melanocytes co-segregate with newly forming scale compartments. Intriguingly, this process coincides with partitioning of both Langerhans cells and dendritic epidermal T-cells to interscale epidermis, suggesting functional segregation of pigmentation and immune surveillance. Analysis of non-pigmented mice and of mice lacking melanocytes or resident immune cells revealed that immunocyte patterning is melanocyte- and melanin-independent, and, *vice versa*, immune cells do not control melanocyte localization. Instead, genetically enforced progressive scale fusion upon *Lrig1* deletion showed that melanocytes and immune cells dynamically follow epithelial scale:interscale patterns. Importantly, disrupting Wnt-Lef1 function in keratinocytes caused melanocyte mislocalization to interscale epidermis, implicating canonical Wnt signaling in organizing the pigmentation pattern. Together, this work uncovered cellular and molecular principles underlying the compartmentalization of tissue functions in skin.

**SUMMARY STATEMENT:** Pigmentation and immune surveillance functions in murine tail skin are spatially segregated by Lrig1- and Wnt-Lef1-dependent keratinocyte lineages that control the partitioning of melanocytes and tissue-resident immune cells into distinct epidermal niches.

## INTRODUCTION

The skin acts as dynamic interface between the organism and its environment. It protects from uncontrolled water loss and external threats such as pathogens, toxins, mechanical damage and temperature variation. Its outermost stratified epidermis is continuous with epidermal appendages like hair follicles and sweat glands (Chuong and Noveen, 1999). Proliferative, undifferentiated basal layer keratinocytes (KCs) attach to the underlying basement membrane, and following asymmetric cell division or delamination progressively differentiate to constitute the spinous, granular and cornified layers (Blanpain and Fuchs, 2006; Dias Gomes and Iden, 2021; Gonzales and Fuchs, 2017; Ray and Lechler, 2011). The epidermis also hosts other resident cell types with crucial functions for the organism: neural-crest derived melanocytes (MCs) provide melanin for hair colorization and to protect KCs from UV damage (D’Orazio et al., 2013), whereas Langerhans cells (LCs) and dendritic epidermal T-cells (DETCs) constitute a first line of defence against environmental pathogens and malignant transformation in murine skin (Deckers et al., 2018; MacLeod et al., 2013). Various UV-induced KC-derived soluble factors have been implicated in melanin induction in MCs (Upadhyay et al., 2021). Yet, how KC, LC, DETC and MC networks are spatiotemporally coordinated within the densely packed epidermis to simultaneously ensure skin barrier, pigmentation and host defence is largely unresolved.

The interfollicular epidermis (IFE) of murine ear and back-skin harbours evenly distributed immune cells (Bergstresser et al., 1980) and is largely devoid of MCs, which mostly reside in the dermis (Hirobe, 1984; Kunisada et al., 2000). In adult ear epidermis, LCs and DETCs actively maintain a non-random distribution that depends on KC density and engages LC- expressed Rac1 (Park et al., 2021), yet the molecular mechanisms how KCs control resident immune cells remain open. Tail skin instead exhibits two distinct epidermal compartments: ‘Scale’ IFE forms a spherical epidermal patch just above hair-follicle triplets and undergoes parakeratotic differentiation, characterized by lack of a granular layer and nucleated cornified layer KCs. ‘Interscale’ IFE surrounds the scale IFE and, alike ear and back-skin IFE, displays orthokeratotic differentiation with a granular layer and enucleated cornified layer KCs (Didierjean et al., 1983). Gomez *et al*. (2013) demonstrated that epidermal lineages of scale and interscale IFE express unique marker genes and are regulated by Wnt, Edaradd, and Lrig1. Strikingly, these intraepidermal patterns in tail skin correlate with a strict compartmentalization of LCs, which populate the interscale IFE (Schweizer and Marks, 1977), and of pigmented MCs, which localize to the scale IFE where they persist throughout adulthood (Glover et al., 2015; Gomez et al., 2013). Further, a small population of amelanotic, quiescent MCs in interscale IFE has been reported (Glover et al., 2015; Köhler et al., 2017). How such segregation of epidermis-resident cells is achieved and maintained is poorly understood. Gomez *et al*. (2013) showed that MCs are dispensable for scale IFE formation; however, molecular and cellular signals orchestrating the mutually exclusive MC:immune cell distribution remain to be identified. It is currently not known whether scale-based MCs antagonize the localization of immune cells (or *vice versa*). Mouse scale IFE closely resembles the epidermal MC distribution of human skin and hence is used to study mechanisms of skin pigmentation (Fitch et al., 2003; Van Raamsdonk et al., 2004) and intraepidermal melanoma (Köhler et al., 2017). Gaining insight into heterologous communication between different epidermis-resident cell types will thus be beneficial to better understand human skin physiology and disease. In this study, we therefore set out to reveal potential interactions and interdependencies of KCs, MCs and immune cells regarding functional compartmentalization in murine tail skin.

## RESULTS

### DETCs, LCs and MCs reside in distinct compartments of the tail IFE

To first map the localization of DETCs, LCs and MCs with respect to IFE niches we performed immunohistochemistry on tail epidermis whole mounts of 6-15 week old C57BL/6 mice. Co-immunostaining for Keratin-31 (K31) marked parakeratotic differentiation and hence scale IFE in epidermal tail whole-mounts (Gomez et al., 2013). In agreement with earlier reports, Trp2/DCT-positive MCs were highly enriched in scale IFE (Glover et al., 2015; Gomez et al., 2013) (Fig. 1A,C,F,I). In the adjacent K31-negative interscale IFE, only few MCs were found (Fig. 1A,C,F), producing melanin and exhibiting a morphology comparable to scale MCs (Fig. S1A-D). Intriguingly, resident immune cells were largely excluded from scale IFE: CD207/Langerin-positive LCs were confined to interscale IFE (Schweizer and Marks, 1977)(Fig. 1A,D,G,I), and co-immunostaining of γδTCR (marking DETCs) and Langerin revealed that DETCs and LCs co-distribute within the interscale IFE (Fig. 1B,E,H,I). Together, these data underpin a remarkable separation of epidermal pigmentary units and resident immune cells in tail IFE, opening the possibility that the scale IFE is an immune-privileged niche.

**Figure 1:**
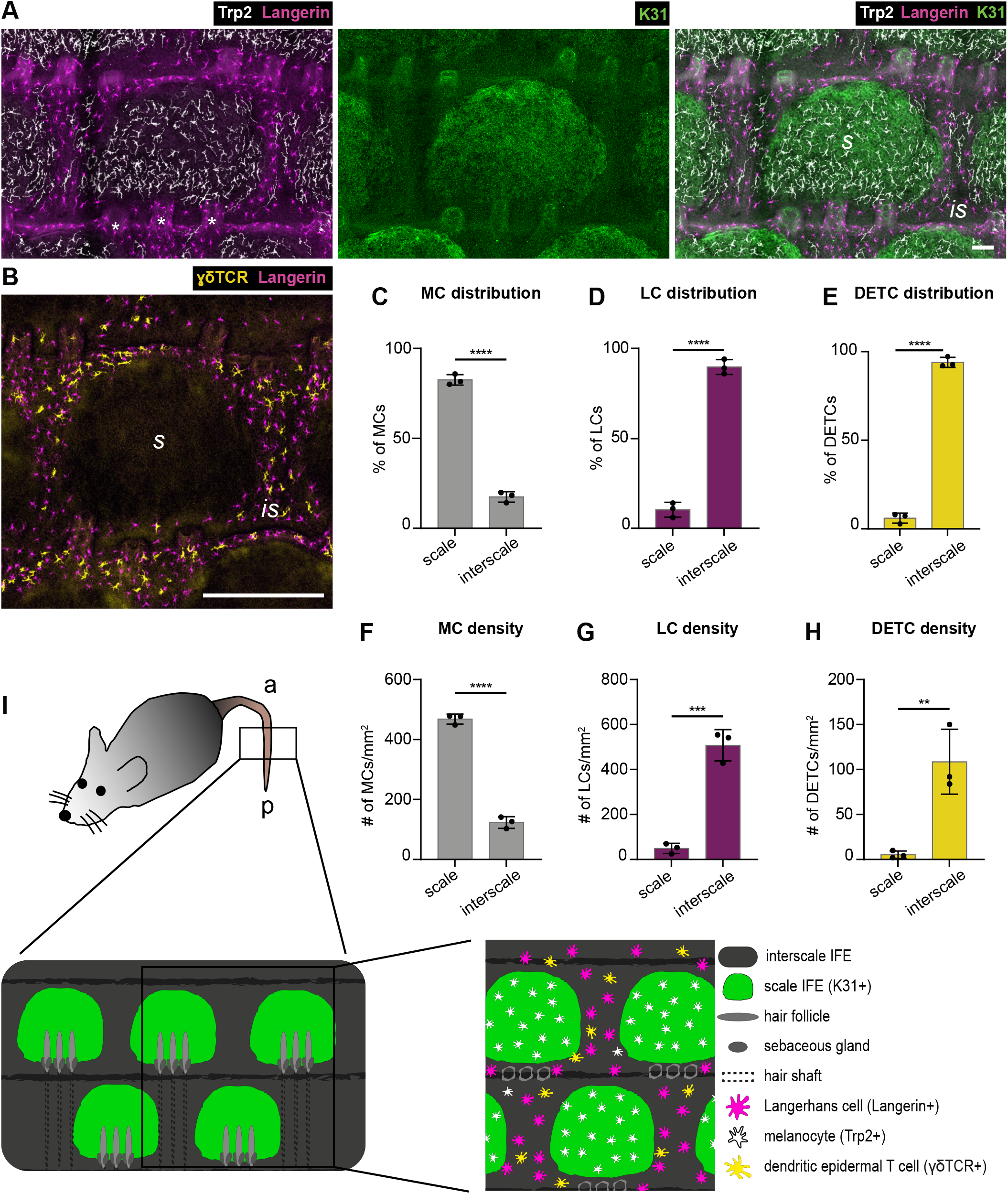
Mutually exclusive localization of MCs and epidermis-resident immunocytes to scale and interscale IFE compartments of murine tail epidermis. (A) Micrographs of tail epidermis whole-mounts from 3-months old wild-type C57BL/6 mice, immunostained for markers of LCs (Langerin), MCs (Trp2/DCT), and scale IFE (K31). Representative for n=4. Scale bar: 75 µm. (B) Immunostaining of tail epidermis whole-mounts from 3-months old wild- type C57BL/6 mice for Langerin and DETC marker γδTCR. Representative for n=3. Scale bar: 300 µm. (C) Quantification of A; MC distribution per scale:interscale unit (% of MCs) in tail epidermis. MC numbers in each compartment (scale, interscale) were normalized to the total MC number per scale:interscale unit. n=3; ****: p<0.0001; mean±s.d.; unpaired Student’s t- test. (D) Quantification of A; LC distribution per scale:interscale unit (% of LCs) in tail epidermis. LC numbers in each compartment (scale, interscale) were normalized to the total LC number per scale:interscale unit. n=3; ****: p<0.0001; mean±s.d.; unpaired Student’s t-test. (E) Quantification of B; DETC distribution per scale:interscale unit (% of DETCs) in tail epidermis. DETC numbers in each compartment (scale, interscale) were normalized to the total DETC number per scale:interscale unit. n=3; ****: p<0.0001; mean±s.d.; unpaired Student’s t-test. (F) Quantification of A; MC density (MC number/mm^2^) in tail epidermis. n=3; ****: p<0.0001; mean±s.d.; unpaired Student’s t-test. (G) Quantification of A; LC density (LC number/mm^2^) in tail epidermis. n=3; ***: p=0.0004; mean±s.d.; unpaired Student’s t-test. (H) Quantification of B; DETC density (DETC number/mm^2^) in tail epidermis. n=3; **: p=0.0079; mean±s.d.; unpaired Student’s t-test. (I) Schematic illustration of scale:interscale IFE patterns and associated structures and cell types in murine tail epidermis; a: anterior, p: posterior, s: scale, is: interscale.

### IFE-resident cell types already segregate during postnatal tail skin development

To shed light into the establishment of these MC:immune cell patterns we next examined early postnatal tissues at the onset of scale formation (Gomez et al., 2013). At postnatal day (P) 5, MCs were already prominently enriched at sites of scale induction (Fig. 2A), a phenomenon that reinforced towards P10 (Fig. 2A). This indicated that MCs co-segregate with the parakeratotic lineage early during skin development. MHCII immunostaining served to visualize LCs at P5 as Langerin is weakly expressed at these stages (Tripp et al., 2004). MHCII, similar to Langerin, is solely found on LCs in unchallenged epidermis, enabling their identification (Tsuruta et al., 1999). Albeit low in numbers due to the postnatal LC proliferation burst yet to occur (Chang-Rodriguez et al., 2005; Kobayashi et al., 1987), LCs were confined to interscale IFE in P5 tail epidermis (Fig. 2A,D). The differential distribution of MCs, LCs and DETCs became even more pronounced with further scale expansion (P10 to P21; Fig. 2A-F), culminating in the dense LC and DETC networks in interscale and MC networks in scale IFE observed in adult tail skin (Fig. 1A-I). These data thus demonstrate that the formation of IFE compartments coincides with the patterned distribution of MCs and epidermis-resident immune cells.

**Figure 2:**
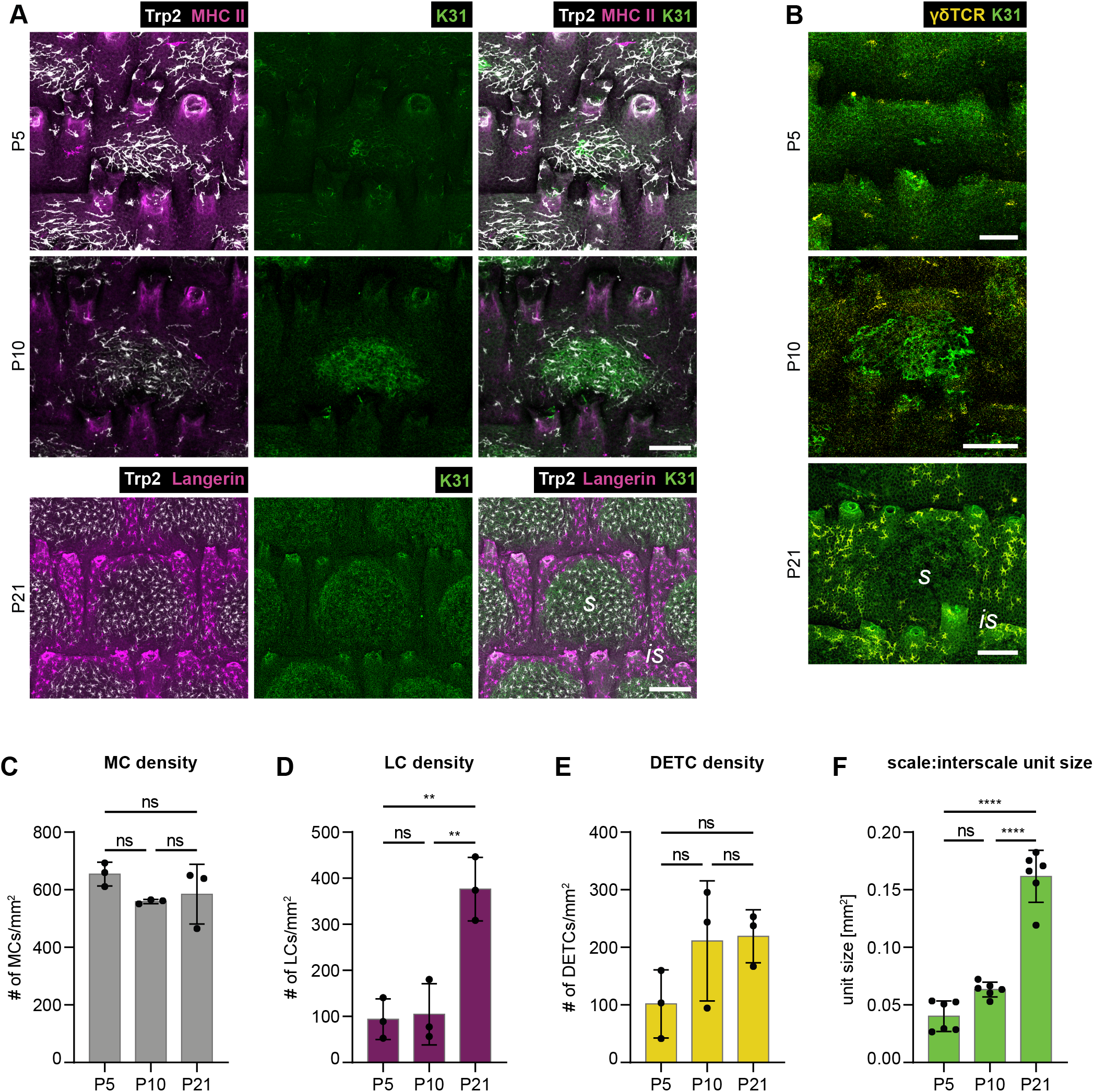
Progressive segregation of IFE-resident cell types during postnatal tail skin development. (A) Immunostaining of tail epidermis whole-mounts from 3-months old wild-type C57BL/6 mice for Trp2, K31, and LC markers MHCII (P5, P10) or Langerin (P21). Representative for n=3. Scale bars: 75 µm. (B) Immunostaining of tail epidermis whole-mounts from 3-months old wild-type C57BL/6 mice for K31 and γδTCR. Representative for n=3. Scale bars: 100 µm. (C) Quantification of A; MC density (MC number/mm^2^) in tail epidermis. n=3; ns: p=0.2458 (P5 vs. P10); ns: p=0.4338 (P5 vs. P21); ns: p=0.8820 (P10 vs. P21); mean±s.d.; one-way ANOVA/Tukey’s multiple comparisons test. (D) Quantification of A; LC density (LC number/mm^2^) in tail epidermis. n=3; ns: p=0.9750 (P5 vs. P10); **: p=0.0031 (P5 vs. P21); **: p=0.0038 (P10 vs. P21); mean±s.d.; one-way ANOVA/Tukey’s multiple comparisons test. (E) Quantification of B; DETC density (DETC number/mm^2^) in tail epidermis. n=3; ns: p=0.2455 (P5 vs. P10); ns: p=0.2082 (P5 vs. P21); ns: p=0.9905 (P10 vs. P21); mean±s.d.; one-way ANOVA/Tukey’s multiple comparisons test. (F) Quantification of A and B; scale:interscale unit size (mm^2^). n=6; ns: p=0.0523 (P5 vs. P10); ****: p<0.0001 (P5 vs. P21); ****: p<0.0001 (P10 vs. P21); mean±s.d.; one-way ANOVA/Tukey’s multiple comparisons test. P: postnatal, s: scale, is: interscale.

### Distinct IFE localization of MCs and immune cells is not interdependent

We next aimed to dissect a potential hierarchy underlying the positioning of MCs, LCs, DETCs and KC lineages. The role of resident immune cells in patterning of the other cell types was analyzed using mice deficient for the transcription factor Id2, which are characterized by complete LC and DETC loss by young adulthood (Hacker et al., 2003; Seré et al., 2012; Yokota et al., 1999). As expected, LCs and DETCs were undetectable in adult *Id2*KO tail epidermis (Fig. 3A,C). The scale:interscale IFE pattern of mutant mice, however, was comparable to control mice (Fig. S2A), and MCs were enriched within scale IFE at the age of 3 months (Fig. 3A,B) up to one year (S.C. Baess, unpublished observation). Likewise, numbers of MCs were similar in control and mutant mice (Fig. S2B). This suggests that both MC distribution and KC lineage progression into scale and interscale identity are independent of resident immune cells. We cannot formally rule out that a transient presence of immune cells during ontogeny might contribute to initial MC and IFE patterning. Nevertheless, our data clearly demonstrate that LCs and DETCs are not required for scale confinement of MCs throughout adulthood.

**Figure 3:**
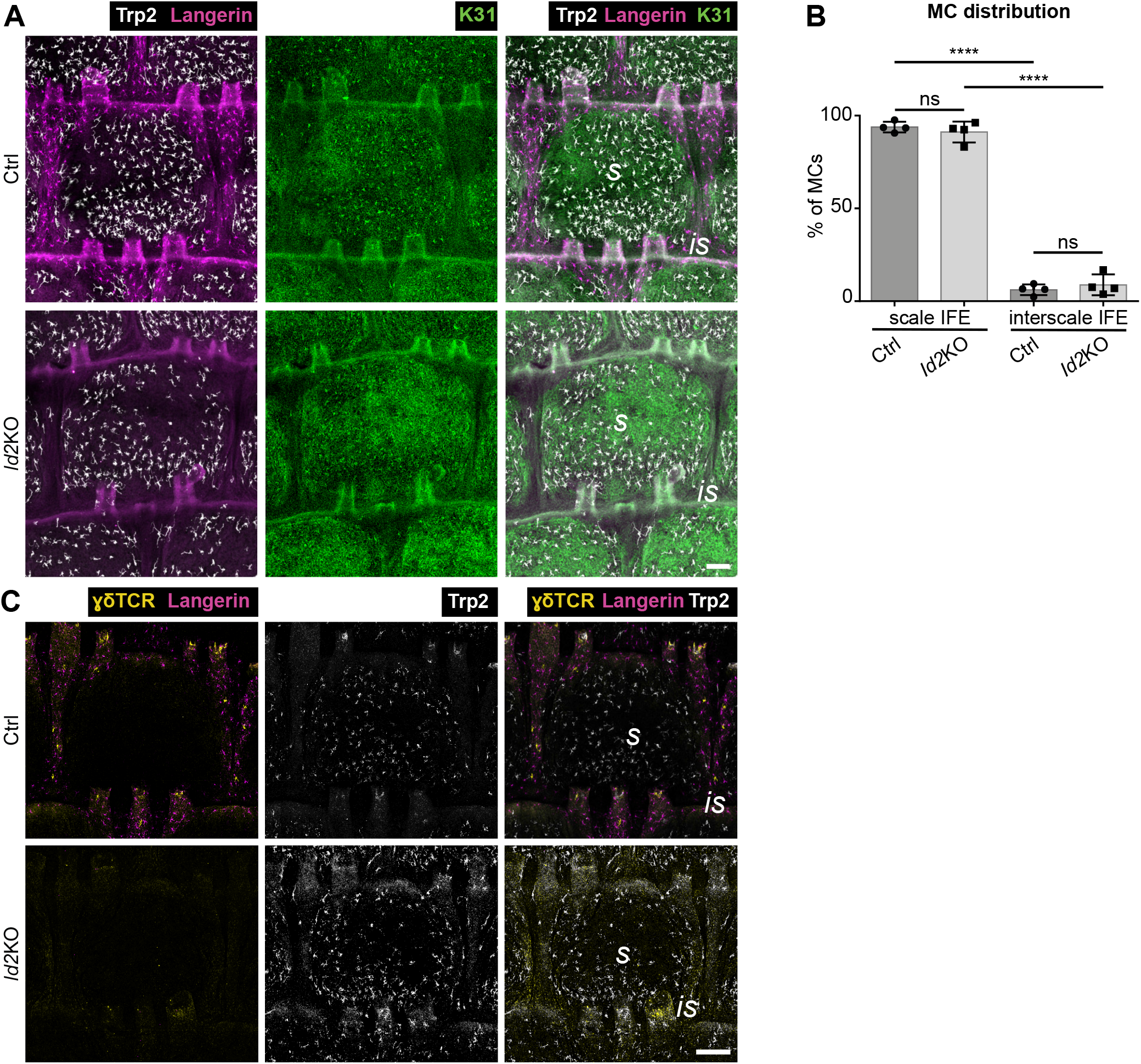
MC localization to scale IFE is independent of epidermis-resident immune cells. (A) Langerin, Trp2 and K31 immunostaining of tail epidermis whole-mounts from 3- months old control and *Id2*KO mice. Representative for n=4. Scale bar: 100 µm. (B) Quantification of A; MC distribution per scale:interscale unit (% of MCs). MC numbers in each compartment (scale, interscale) were normalized to the total MC number per scale:interscale unit. n=4; ns: p=0.8370; ****: p<0.0001; mean±s.d.; one-way ANOVA/Tukey’s multiple comparisons test. (C) Langerin and γδTCR immunostaining of tail epidermis whole-mounts from 3-months old control and *Id2*KO mice. Representative for n=3. Scale bar: 300 µm. s: scale, is: interscale

Next we asked if, *vice versa*, MCs antagonize the localization of IFE-resident immune cells. Insects use melanin as innate immune defense (Gillespie et al., 1997), and immunomodulatory roles have also been proposed for mammalian MCs and MC-derived melanin (Burkhart and Burkhart, 2005; Hong et al., 2015). LCs can take up melanin (Breathnach and Wyllie, 1965; Mishima, 1966; Tobin, 1998), which may cause LC emigration to draining lymph nodes (Hemmi et al., 2001). We therefore explored if MCs, or melanin levels, counteract immune cell localization to scale IFE. Comparison of adult pigmented C57BL/6 mice and non-pigmented (albino) FVB/N mice, however, revealed no difference of MC, LC and DETC distribution per scale:interscale unit (Fig. 4A-E). Similarly, localized spontaneous depletion of MCs and subsequent loss of pigmentation in tail epidermis of C57BL/6 mice (Fig. 4F) did not cause redistribution of LCs or DETCs into the MC-free scale IFE (Fig. 4G-J). Of note, despite stable immune cell distributions, DETC densities were reduced in FVB/N mice but not in mice with spontaneous MC loss (Fig. S3A-E), pointing to strain-dependent variation in numbers of DETCs. Together, epidermal MCs and resident immune cells do not affect each other’s positioning in tail epidermis.

**Figure 4:**
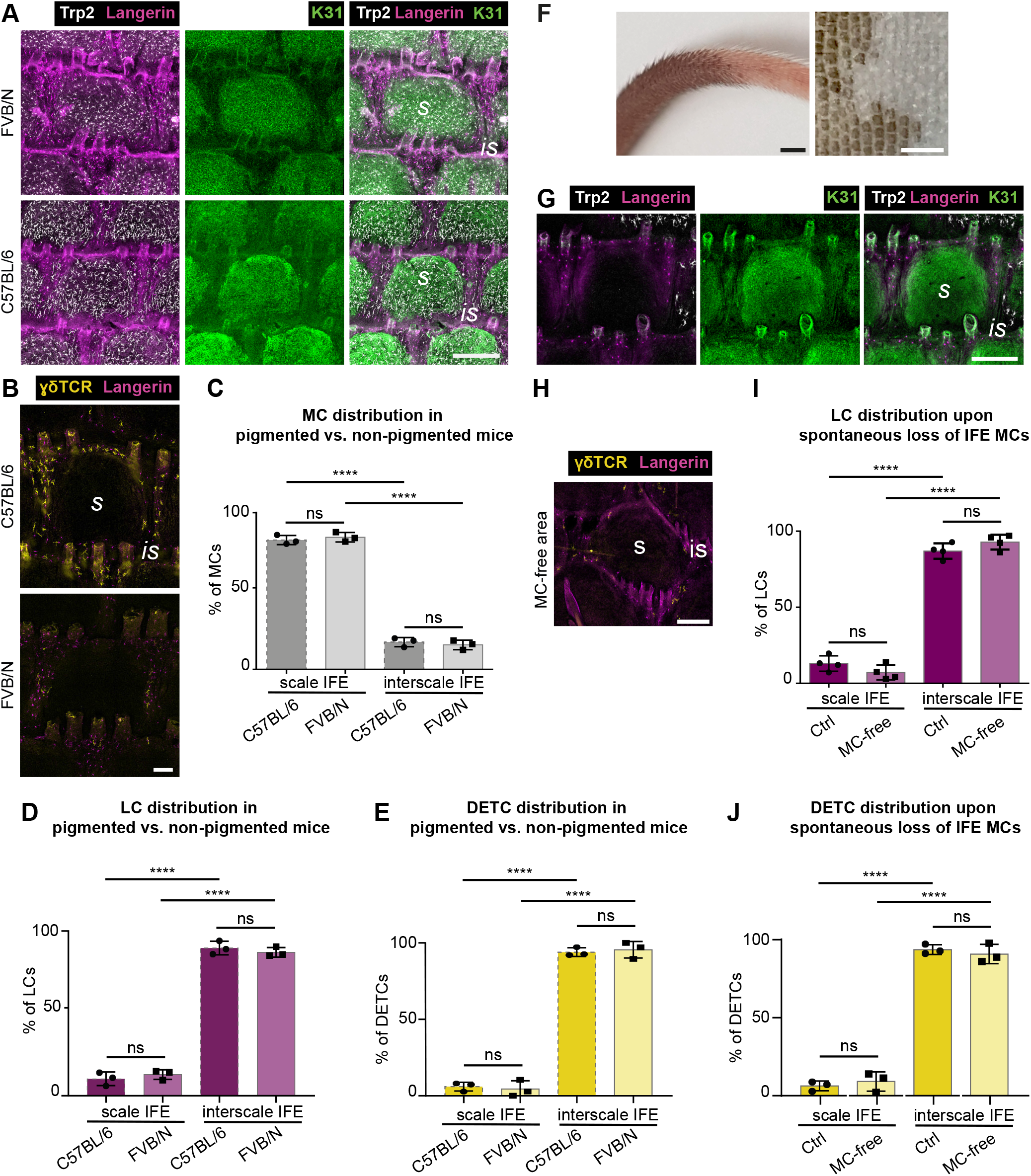
LCs and DETCs localize independently of MCs and melanin production to interscale IFE. (A) Langerin, Trp2 and K31 immunostaining of tail epidermis whole-mounts of 3-months old wild-type C57BL/6 and FVB/N mice. Representative for n=3. Scale bar: 250 µm. (B) γδTCR and Langerin immunostaining of tail epidermis whole-mounts of 3-months old wild- type C57BL/6 and FVB/N mice. Representative for n=3. Scale bar: 100 µm. (C) Quantification of A; MC distribution to scale and interscale IFE (% of MCs). MC numbers in each compartment (scale, interscale) were normalized to the total MC number per scale:interscale unit. n=3; ns: p=0.8755; ****: p<0.0001; mean±s.d.; one-way ANOVA/Tukey’s multiple comparisons test. Data for C57BL/6 as shown in Figure 1C. (D) Quantification of A; LC distribution per scale:interscale unit (% of LCs). LC numbers in each compartment (scale, interscale) were normalized to the total LC number per scale:interscale unit. n=3; ns: p=0.8186; ****: p<0.0001; mean±s.d.; one-way ANOVA/Tukey’s multiple comparisons test. Data for C57BL/6 as shown in Fig. 1D. (E) Quantification of B; DETC distribution per scale:interscale unit (% of DETCs). DETC numbers in each compartment (scale, interscale) were normalized to the total DETC number per scale:interscale unit. n=3; ns: p=0.9661; ****: p<0.0001; mean±s.d.; one-way ANOVA/Tukey’s multiple comparisons test. Data for C57BL/6 as shown in Fig. 1E. (F) Representative images of tail skin from 3-months old wild-type C57BL/6 with partial spontaneous loss of MCs. Left: intact tail skin. Right: epidermis whole-mount. Representative for n=4. Scale bars: 3000 µm (left), 1500 µm (right). (G) Langerin, Trp2 and K31 immunostaining of tail epidermis whole-mounts of 3-months old wild-type C57BL/6 mice with spontaneous loss of MCs (center) in posterior tail region. Representative for n=4. Scale bar: 250 µm. (H) Langerin and γδTCR immunostaining of tail epidermis whole-mounts of 3-months old wild-type C57BL/6 mice with spontaneous loss of MCs in posterior tail epidermis. Representative for n=3. Scale bar: 150 µm. (I) Quantification of G; LC distribution per scale:interscale unit (% of LCs) in MC-free and MC-containing tail epidermis. LC numbers in each compartment (scale, interscale) were normalized to the total LC number per scale:interscale unit. n=4; ns: p=0.3839; ****: p<0.0001; mean±s.d.; one-way ANOVA/Tukey’s multiple comparisons test. (J) Quantification of H; DETC distribution per scale:interscale unit (% of DETCs) in MC-free and MC-containing tail epidermis. DETC numbers in each compartment (scale, interscale) were normalized to the total DETC number per scale:interscale unit. n=3; ns: p=0.8854; ****: p<0.0001; mean±s.d.; one-way ANOVA/Tukey’s multiple comparisons test. s: scale, is: interscale.

### MC:immune cell distribution follows scale:interscale patterning

Our observations and the strikingly similar dynamics of scale formation and MC clustering during postnatal development (Fig. 2A) prompted us to investigate the role of parakeratotic scale and orthokeratotic interscale IFE compartments for immunocyte and MC patterning. Scale progenitors divide more rapidly than interscale progenitors (Gomez et al., 2013; Sada et al., 2016), with spatial control of scale progenitor proliferation likely contributing to scale:interscale patterning and boundary formation. Lrig1, a negative regulator of EGFR signaling, is expressed in the dermis underneath interscale IFE where it is thought to antagonize scale fate and size through dermal-epidermal signaling (Gomez et al., 2013). Upon constitutive *Lrig1* deletion, initial scale induction is normal but scale IFE compartments progressively enlarge and eventually laterally fuse into wide dorsoventral bands in adult mice (Gomez et al., 2013)(Fig. 5A). Interestingly, analysis of *Lrig1*KO mice at P10 and adult stages unraveled that, concomitant with lateral scale fusion, MCs assumed a band-like expansion corresponding to the K31-labelled scale band, whereas LCs and DETCs were restricted to the diminished interscale stripe (Fig. 5A-F). Notably, despite their different scale and interscale architectures, both MC and LC densities were comparable in control and *Lrig1*KO mice (Fig. S4A,B). Likewise, LC densities were similar during and after scale fusion in *Lrig1*KO mice (Fig. S4C). This opens the possibility that LCs are expelled or lost when scales fuse; however, we did not observe apoptotic LCs in *Lrig1*KO tail skin (Fig. S4D). Together, above data indicate that patterning of MCs and immune cells is primarily determined by compartmentalization of the epidermis into scale and interscale IFE.

**Figure 5:**
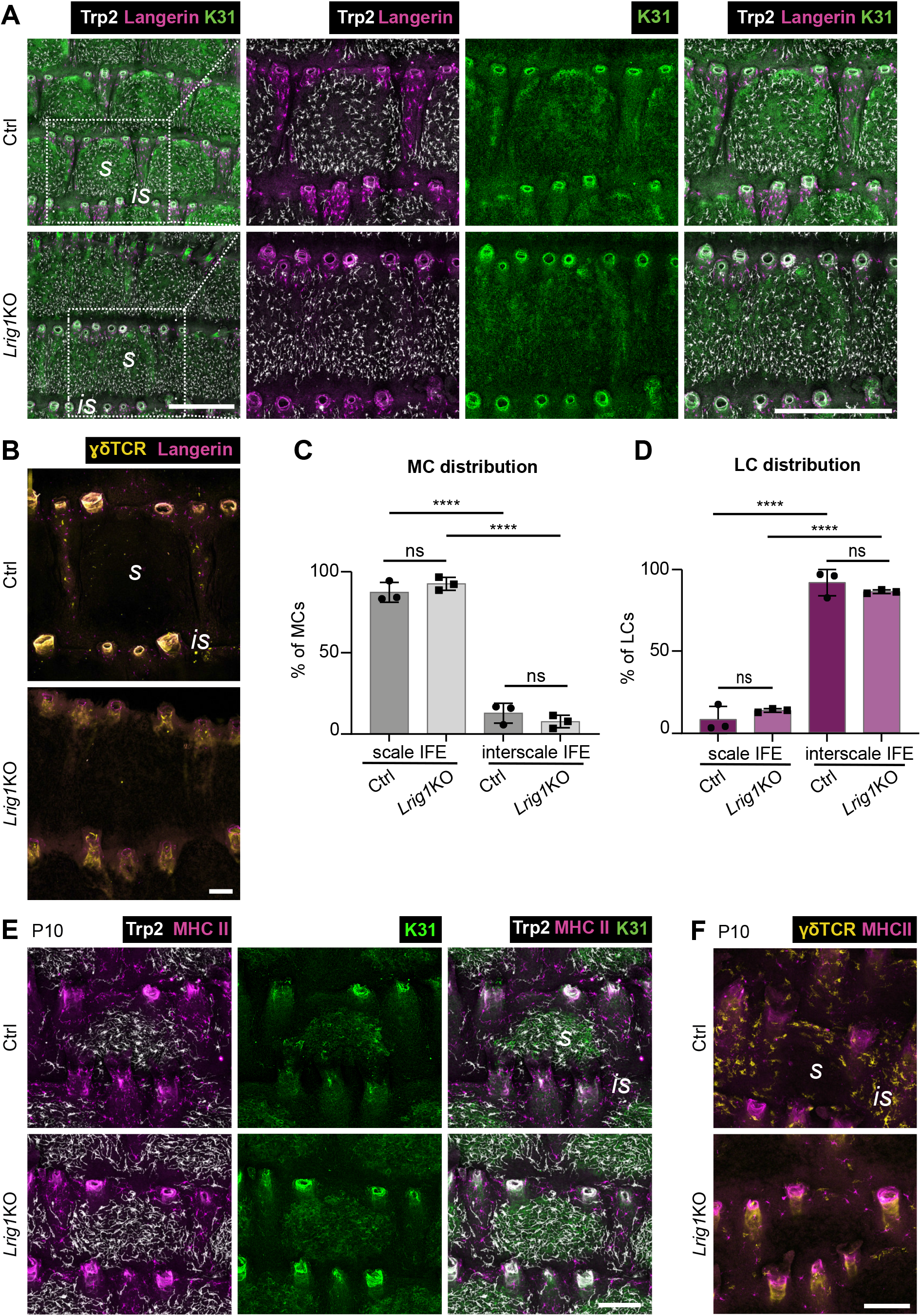
MC:immunocyte distribution dynamically adapts to changes of epidermal scale:interscale patterning in *Lrig1*KO mice. (A) Langerin, Trp2 and K31 immunostaining of tail epidermis whole-mounts from 3-months old control and *Lrig1*KO mice. Representative for n=3. Scale bars: 500 µm. (B) γδTCR and Langerin immunostaining of tail epidermis whole- mounts from 3-months old control and *Lrig1*KO mice. Representative for n=3. Scale bar: 100 µm. (C) Quantification of A; MC distribution per scale:interscale unit (% of MCs). MC numbers in each compartment (scale, interscale) were normalized to the total MC number per scale:interscale unit. n=3; ns: p=0.6203; ****: p<0.0001; mean±s.d.; one-way ANOVA/Tukey’s multiple comparisons test. (D) Quantification of A; LC distribution per scale:interscale unit (% of LCs). LC numbers in each compartment (scale, interscale) were normalized to the total LC number per scale:interscale unit. n=3; ns: p=0.6504; ****: p<0.0001; mean±s.d.; one-way ANOVA/Tukey’s multiple comparisons test. (E) MHCII, Trp2 and K31 immunostaining of tail epidermis whole-mounts from control and *Lrig1*KO mice at P10. Representative for n=4. Scale bar: 100 µm. (F) MHCII and γδTCR immunostaining of tail epidermis whole-mounts from control and *Lrig1*KOmice at P10. Representative for n=4. Scale bar: 100 µm. P: postnatal, s: scale, is: interscale.

### Defective epidermal Wnt-Lef1 signaling causes MC redistribution to interscale IFE

We next set out to identify intraepidermal factors mediating MC and immune cell patterning. Among candidate pathways, we hypothesized that canonical Wnt signaling could be involved as it is known for steering epidermal lineage selection in postnatal epidermis (Watt and Collins, 2008) and because Lef1, a key transcriptional effector of canonical Wnt signaling, is predominantly expressed in scale IFE (Gomez et al., 2013; Niemann et al., 2002). Expression of N-terminally truncated Lef1 in epidermal KCs (K14ΔNLef1; Niemann et al., 2002), acting as dominant negative inhibitor of canonical Wnt signaling, results in impaired scale differentiation and size with irregular scale:interscale patterning (Gomez et al., 2013). Strikingly, analysis of skin-resident cell types in these mice revealed that MC distribution was largely inverted compared to control mice, with MCs localizing to interscale IFE and to the scale periphery, whereas the scale center was mostly devoid of MCs (Fig. 6A-D). Next to this repositioning, total numbers of MCs in K14ΔNLef1 tissues were reduced (Fig. S5A,B), and remaining scale-based MCs showed increased mean area, axis length and dendricity (Fig. S6A-F). Interscale MCs in mutant mice retained the ability to produce melanin (Fig. S6G) and did not show signs of increased apoptosis (Fig. S6H). Congruent with the redistribution of MCs in mutant tail skin, MC numbers were significantly increased in K14ΔNLef1 ear epidermis while rarely detected in control ear epidermis (Fig. 6E,F). These combined findings in tail and ear epidermis strongly suggest that inhibition of canonical Wnt signaling renders orthokeratotic epidermis permissive for MC colonization. Notably, resident immune cells remained confined to the interscale IFE in K14ΔNLef1 mice (Fig. 6A,G,H), whereby densities of LCs in interscale regions and of DETCs in ear epidermis of mutant mice were reduced (Fig. S5C-H).

**Figure 6:**
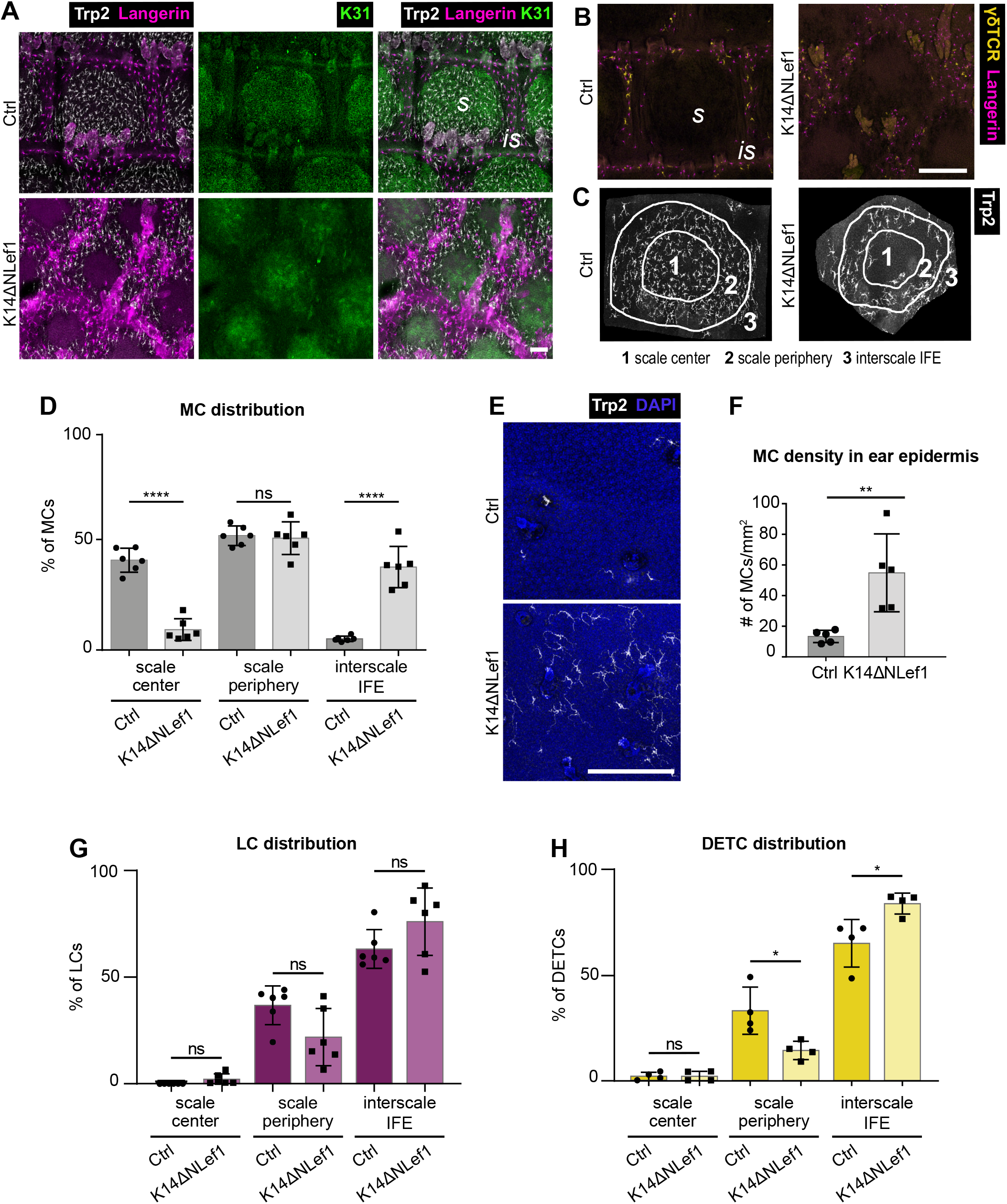
Repression of epidermal Wnt signaling causes ectopic localization of MCs to interscale IFE. (A) Immunostaining of tail epidermis whole-mounts of 3-months old control and K14ΔNLef1 mice for Langerin, Trp2, and K31. Representative for n=6. Scale bar: 100 µm. (B) γδTCR and Langerin immunostaining of tail epidermis whole-mounts from 3-months old control and K14ΔNLef1 mice. Representative for n=4. Scale bar: 250 µm. (C) Illustration of one scale:interscale unit with corresponding categorization into scale center, scale periphery, and interscale IFE for analysis of MC distribution in K14ΔNLef1 mice as done in D, G and H. (D) Quantification of A; MC distribution per scale:interscale unit (% of MCs) in control and K14ΔNLef1 mice. MC numbers per compartment were normalized to the total number of the scale:interscale unit. n=6; ****: p<0.0001 (scale center, Ctrl vs. K14ΔNLef1); ns: p=0.9995 (scale periphery, Ctrl vs. K14ΔNLef1); ****: p<0.0001 (interscale IFE, Ctrl vs. K14ΔNLef1); mean±s.d.; one-way ANOVA/Tukey’s multiple comparisons test. (E) Trp2 immunostaining of ear epidermis whole-mounts of 3-months old control and K14ΔNLef1 mice. Nuclei were counterstained using DAPI. Representative for n=5. Scale bar: 250 µm. (F) Quantification of E; MC density (MC number/mm^2^) in ear epidermis. n=5; **: p=0.0079; mean±s.d.; Mann- Whitney test. (G) Quantification of A; LC distribution per scale:interscale unit (% of LCs) in control and K14ΔNLef1 mice. LC numbers per compartment were normalized to the total number of the scale:interscale unit. n=6; ns: p=0.9991 (scale center, Ctrl vs. K14ΔNLef1); ns: p=0.1341 (scale periphery, Ctrl vs. K14ΔNLef1); ns: p=0.2596 (interscale IFE, Ctrl vs. K14ΔNLef1); mean±s.d.; one-way ANOVA/Tukey’s multiple comparisons test. (H) Quantification of B; DETC distribution per scale:interscale unit (% of DETCs) in control and K14ΔNLef1 mice. DETC numbers per compartment were normalized to the total number of the scale:interscale unit. n=6; ns: p>0.9999 (scale center, Ctrl vs. K14ΔNLef1); *: p=0.0163 (scale periphery, Ctrl vs. K14ΔNLef1); *: p=0.0166 (interscale IFE, Ctrl vs. K14ΔNLef1); mean±s.d.; one-way ANOVA/Tukey’s multiple comparisons test. s: scale, is: interscale.

To understand if the observed striking MC mislocalization upon altered Wnt-Lef1 function is a progressive event, we next assessed the distribution of resident cell types in mutant tissues during postnatal development. K31 immunolabelling of P5 and P10 tail epidermis illustrated that scale formation in K14ΔNLef1 mice was delayed when compared to control mice (Fig. 7A)(Gomez et al., 2013). Importantly, already at this early stage MCs failed to cluster to scale IFE and instead overlapped with LCs in the K31-negative compartment in K14ΔNLef1 mice early-on (Fig. 7A,B). This indicates that canonical Wnt signaling in KCs orchestrates not only scale formation and maintenance but also the distinct patterning of epidermis-resident MCs and hence tail skin pigmentation.

**Figure 7:**
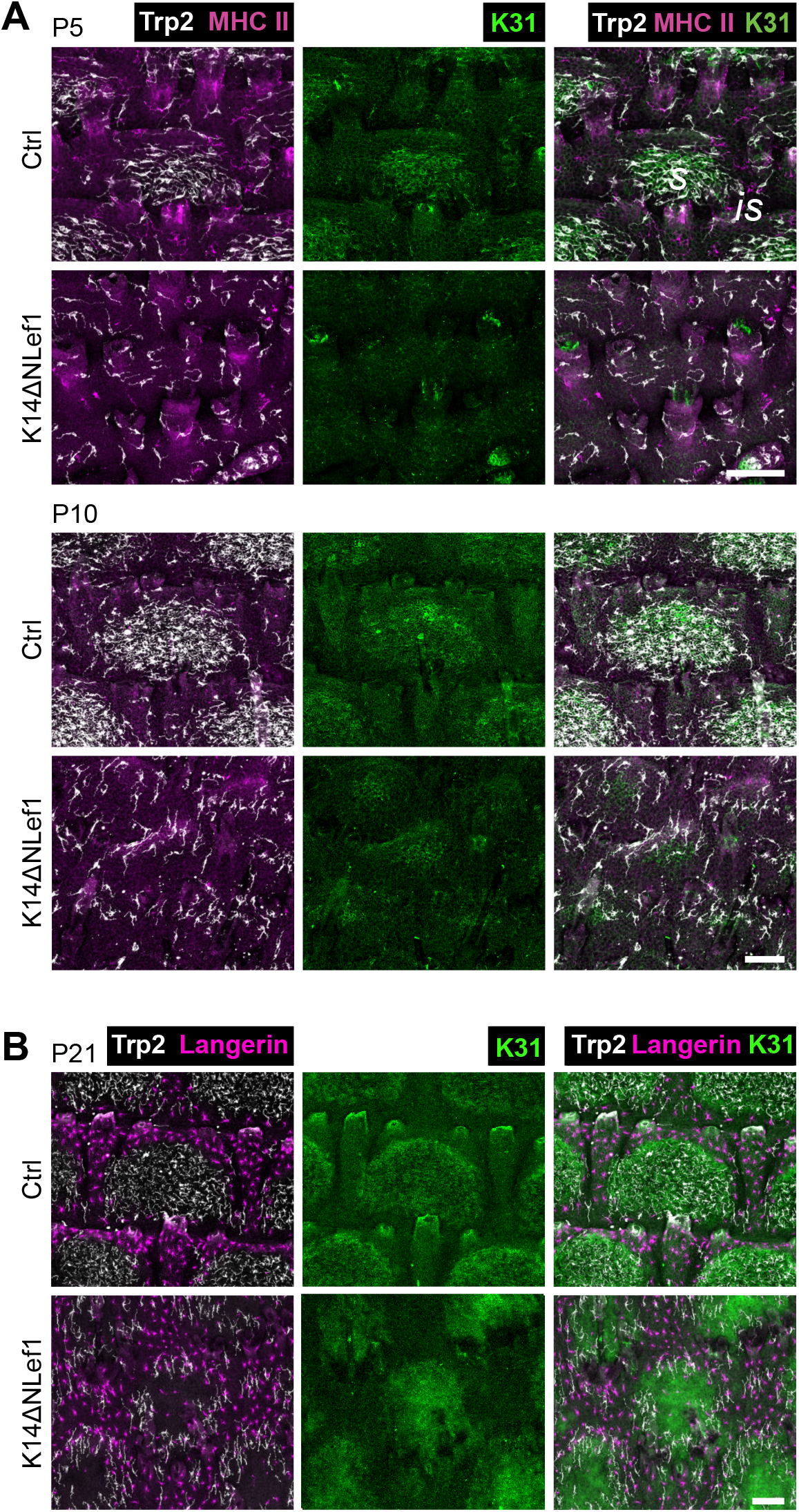
During postnatal tail skin development, MCs in K14ΔNLef1 mice fail to enrich in forming scale regions. (A) Trp2, K31, and MHCII immunostaining of tail epidermis whole- mounts of control and K14ΔNLef1 mice at P5 and P10. Representative for n=3. Scale bars: 100 µm. (B) Trp2, K31, and Langerin immunostaining of tail epidermis whole-mounts from P21 control and K14ΔNLef1 mice. Representative for n=3. Scale bar: 100 µm. P: postnatal, s: scale, is: interscale.

Collectively, these findings demonstrate that the mutually exclusive localization of MCs and epidermis-resident immune cells critically depends on epidermal patterns that are orchestrated by antagonistic signaling downstream of Lrig1 and Wnt-Lef1 (Fig. 8A). Our work identified epidermal KCs at the top of a cellular hierarchy that guides the formation and maintenance of functionally distinct tail IFE compartments.

**Figure 8:**
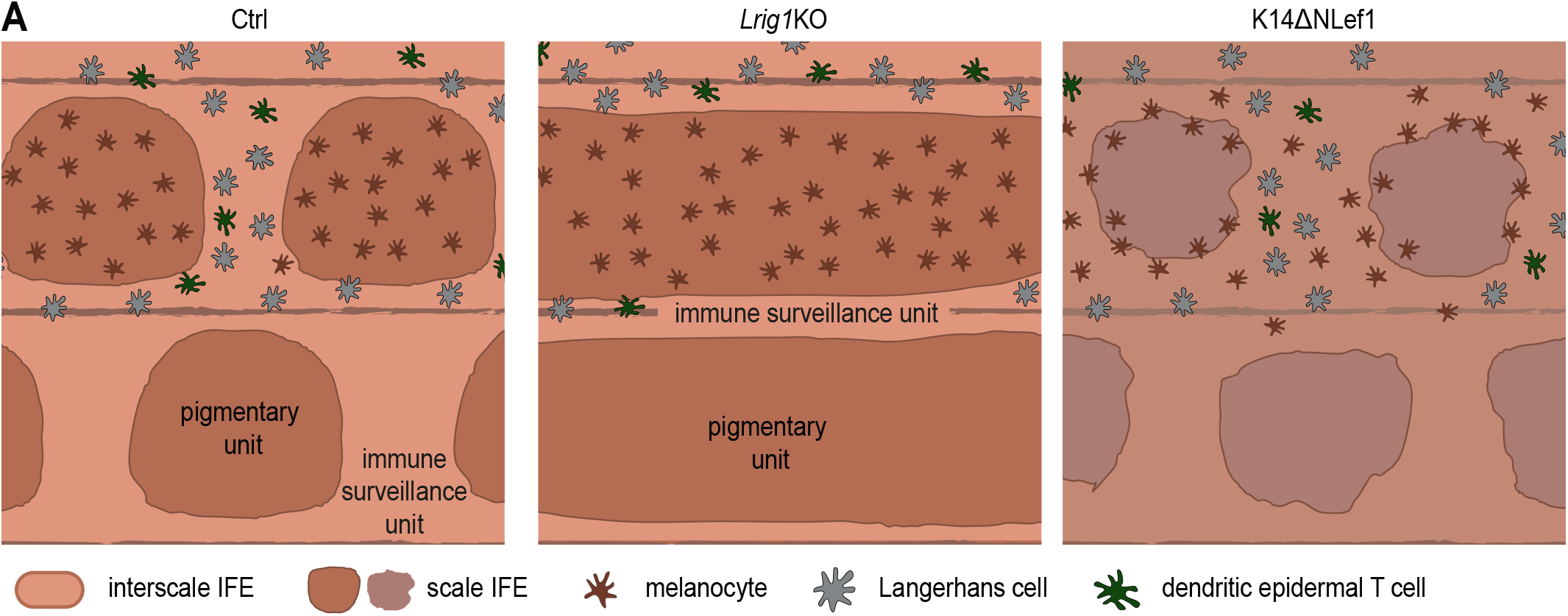
Schematic summary of Lrig1- and Wnt-dependent partitioning of MCs and resident immunocytes into distinct epidermal niches. **(A)** Epidermal niches and their resident cell types in control mice (left), *Lrig1*KO mice exhibiting scale IFE fusion (middle), and K14ΔNLef1 mice with MC localization inverted to interscale IFE, resulting in loss of functional segregation of pigmentary units and immune surveillance unit in mouse tail skin (right panel).

## DISCUSSION

Evolutionarily, pigmentation of scale IFE may have prevailed to protect the underlying hair follicle stem cells from UV-mediated damage. Yet, it is intriguing that immune cells are strictly excluded from this compartment. Melanin synthesis involves generation of cytotoxic intermediates, increasing oxidative stress (Denat et al., 2014; Graham et al., 1978; Urabe et al., 1994); however, we show that the epidermal lineages, but not MCs or melanin, determine LC/DETC patterning. KC-derived paracrine factors (e.g., α-melanocyte stimulating hormone, endothelin, TGFβ) regulate follicular MC stem cell maintenance (Nishimura, 2011) and induction of melanin synthesis (Li et al., 2020; Upadhyay et al., 2021). In contrast, principles of KC:immune cell communication are much less understood. mRNA transfer from KCs to MCs through exosomes (Cicero et al., 2015; Liu et al., 2019) and from LCs to KCs and DETCs has been reported, likely through tunneling nanotubes (Su and Igyártó, 2019), opening the possibility that heterologous cell types in the epidermis actively communicate via such structures. Yet, these studies utilized *in vitro* systems or non-patterned orthokeratotic epidermis, which leaves open the question of which molecules and signaling pathways mediate the partitioning of epidermis-resident cells to different epidermal niches. Our work identifies an antagonistic framework in which Lrig1-dependent IFE lineage commitment instructs the segregation of epidermal MCs, LC and DETCs, and where epidermal Wnt-Lef1 signaling restricts mature MCs to scale IFE.

Our data derived from the *Lrig1*-KO model, characterized by progressive scale fusion, point to remarkably stable LC densities during interscale shrinkage and hence adaptation of the LC network to changes in scale:interscale IFE ratios. Though not directly comparable, this partly resembles the recently reported modulation of LC numbers upon alterations of KC densities in adult, non-patterned orthokeratotic ear epidermis (Park et al., 2021). These combined findings illustrate previously unrecognized dynamics of resident immune cells and their ability to adapt to changes in tissue epithelia. Regarding our results on Wnt-Lef1-mediated tissue patterning, further work is necessary to reveal Lef1 transcriptional targets that control MC scale localization. Albeit hypothetical at this point, both the strong decline and the associated morphological alterations of scale-restricted MCs in K14ΔNLef1 IFE shown here, combined with the reported high expression of endogenous Lef1 in wild-type scale IFE (Gomez et al., 2013; Niemann et al., 2002), suggests that canonical Wnt signaling elicits expression of MC- retention factors in the scale IFE. However, as transgene expression in the K14ΔNLef1 model is not restricted to scale IFE, an active recruitment of MCs to K14ΔNLef1 interscale IFE through yet unknown mechanisms is possible as well.

We were also intrigued by previous reports of differential transformation susceptibility of scale and interscale IFE cells: Oncogenic hedgehog signaling causes basal cell carcinoma predominantly in interscale IFE (Sánchez-Danés et al., 2016), whereas oncogenic Braf-induced melanomas arise from pigmented MCs in scale IFE (Köhler et al., 2017). This poses the question whether such differential oncogenic outcome is, next to cell-intrinsic factors like proliferative capacity, also linked to the functional separation of pigmentation and skin immunity. LCs sample environmental antigens (Cho et al., 2010; de Jong et al., 2010; Leclercq and Plum, 2021), present self-antigens to naïve T cells to induce tolerance (Stoitzner et al., 2006), and detect altered self-antigens or neo-antigens following external stresses or during malignant transformation (Cao et al., 2007; Schreurs et al., 2000). The role of LCs in various skin cancers and wound healing, however, remains controversial, likely due to recently identified additional dermal Langerin-positive non-LC populations that require reinterpretation of data from genetic models with langerin promoters (Deckers et al., 2018; Sheng et al., 2021; Li et al., 2021). DETCs instead have been clearly implicated in immune surveillance, shown to protect against skin tumors and to promote wound healing (Girardi et al., 2003; Havran and Jameson, 2010; Jameson et al., 2002; Kaminski et al., 1993; Schuhmachers et al., 1995; Xiang et al., 2020). Albeit unclear whether above immune cell functions also apply to tail skin, this opens the possibility that immune surveillance in interscale IFE, and lack thereof in scale IFE, is decisive for tumorigenesis. Interestingly, MCs themselves may serve as non-professional antigen presenting cells (Le Poole et al., 1993a), perform phagocytosis (Le Poole et al., 1993b) and produce immune-regulatory cytokines and chemokines (Hong et al., 2015). Though requiring further investigation, considering that scale IFE lacks resident immune cells, it seems possible that MCs, in principle, compensate for certain immune cell functions in this compartment. Moreover, it will be interesting to learn whether the rare, pigmented interscale MCs described in this study serve different functions than their scale counterparts, albeit phenotypically comparable (Fig. S1A-D). Additionally, how this pool of MCs relates to the amelanotic interscale MCs reported by Glover *et al*. (2015) and Köhler *et al*. (2017) requires further investigation. Together, dissecting the cellular complexity of epidermal niches and communication of the different resident cell types is an important future task to advance our understanding of cell transformation and malignant progression in the skin.

## MATERIALS AND METHODS

### Mice

All animal breeding and tissue analyses were performed according to institutional guidelines and in compliance with the German federal law for animal protection under control of the North Rhine-Westphalian State Agency for Nature, Environment and Consumer Protection (LANUV, NRW, Germany; file references 81-02.04.2018.A384 and 81-02.04.2018.A401) and the Veterinary Office City of Cologne, Germany (file references UniKöln_Anzeige §4.17.018 and §4.18.020). *Id2*KO (Id2^tm1Yyk^)(Yokota et al., 1999) and K14ΔNLef1 (Niemann et al., 2002) mice have been previously described. *Lrig1*KO mice were generated by crossing Lrig1- CreERT2/EGFP mice (Page et al., 2013) (kindly provided by Prof. Kim B. Jensen, University of Copenhagen, Danstem, Denmark) on 129 background to homozygosity, resulting in *Lrig1* deletion.

### Immunohistochemistry of tail epidermal whole-mounts

Preparation and staining of mouse tail whole-mounts was performed as previously described (Braun, 2003). Briefly, mouse tail skin was peeled off the bone and incubated in 5 mM EDTA/PBS for 2 h at RT with mild shaking at 400 rpm. After replacing half of the EDTA/PBS volume, the tissue was incubated for another 2 h. Next, the epidermis was separated from the dermis leaving hair follicles attached to the epidermal sheet. Depilation was performed to eliminate autofluorescence caused by hair. The epidermis was fixed in pre-cooled acetone for 30 min on ice. Samples stained with directly conjugated antibodies (γδTCR-FITC) were first incubated with the antibody diluted in PBS over-night at 4°C, followed by blocking the next day. For blocking, samples were incubated in PB buffer (0.5% skim milk powder, 0.25% fish skin gelatin (Sigma-Aldrich), 0.5% Triton-X100 in HBS (20 mM HEPES pH 7.2, 0.9% NaCl)) or 3% milk/PBS for 1 h at RT followed by incubation with primary antibodies diluted in blocking buffer over-night at RT. The next day samples were extensively washed for 6-8 h with 0.5% Triton X- 100/PBS and multiple exchanges of washing buffer. Samples were then incubated with secondary antibodies (diluted in blocking buffer) over-night at RT. Finally, washing steps for 6- 8 h were repeated at RT and samples were mounted in Mowiol with the basal side facing upwards.

### Co-immunostaining of K31 and Trp2 and of γδTCR, CD207 and Trp2 in tail epidermal whole-mounts

Due to limited choices of primary antibodies suited for tail skin whole-mount preparations, co-detection of K31 and Trp2 required sequential immunostaining to avoid cross-reactivity of secondary antibodies (AF647 donkey-anti-goat and AF488 goat-anti-guinea pig, see Table S1). Samples were incubated in secondary antibodies except for AF488-conjugated anti-guinea pig over-night at RT, followed by 6-8 h of washing in 0.5% Triton X-100/PBS and subsequent incubation with AF488 goat anti-guinea pig antibodies over-night. Using this protocol, cross-reactivity with Trp2 primary antibodies could be strongly diminished though not completely eliminated. In some samples, dim signals for MCs were noted when acquiring K31 signals. This was attributed to secondary antibodies, as we could validate that the anti-K31 antibody itself does not recognize MC antigens (S.C. Baess, unpublished observation). Importantly, K31 signals solely served to identify scale areas, but not to quantify signal intensities or cell morphologies, hence this weak MC signal was not relevant for the analyses presented in this study.

Similarly, co-detection of γδTCR, CD207 and Trp2 required sequential immunostaining to avoid cross-reactivity of secondary antibodies (AF568 donkey-anti-goat and AF647 goat-anti-rat, see Table S1). Samples were incubated in secondary antibodies except for AF647 goat-anti-rat over-night at RT, followed by 6-8 h of washing in 0.5% Triton X-100/PBS and subsequent incubation with AF647 goat-anti-rat antibodies over-night.

### Immunohistochemistry of ear epidermal whole-mounts

Preparation and staining of mouse ear epidermis whole-mounts was performed as previously described (Ross et al., 1998). Briefly, mouse ears were depilated with Veet Hair Removal Cream and residual ventral cartilage was removed. Ears were split while floating on PBS, and for further use only the inner ear skin was kept. To separate dermis and epidermis, the skin was placed on 0.5 M ammonium thiocyanate for 25 min at 37°C with the dermal side down. After removing the dermis from the epidermis using forceps, the epidermis was spread on the bottom of a 12-well plate, covered with ice-cold acetone and fixed on ice for 20 min. Next, ear tissues were blocked in 3% milk/PBS for 2 h at RT and subsequently incubated with primary antibodies over-night at 4°C. The next day, ear epidermis was washed three times in PBS for 20 min at RT, followed by incubation with secondary antibodies (diluted in AB buffer, 0.4% BSA, 0.5% Triton-X100 in PBS) for 2 h at 37°C. Finally, the ear tissues were washed three times in PBS at RT and then mounted in Mowiol, with the basal side facing upwards.

### Melanin detection in tail skin using Fontana-Masson staining

Fontana-Masson silver staining served to visualize melanin in skin tissues, based on melanin-mediated reduction of silver nitrate to metallic silver. For this, the Fontana-Masson Staining Kit (Sigma Aldrich, HT200) was used. Tail epidermal whole-mounts were fixed with 4% PFA/PBS for 30 min at RT and incubated in freshly prepared ammoniacal silver solution at 60°C for 1 min. After rinsing in water, samples were quickly mounted in Mowiol and micrographs directly acquired.

### Microscopy

Confocal images were acquired using a Leica SP8 confocal laser scanning microscope and LASX software using a PL Apo 10x/0.40 CS2 air objective or a Zeiss LSM880 confocal laser scanning microscope and ZEN 2.3 SP1 software using a PL Apo 40x/1.3 Oil DIC UV-IR M27 objective. For detection of Fontana-Masson staining, the 633 nm laser and T-PMT detector were used.

### Analysis of MC, LC and DETC distributions in tail epidermal whole-mounts

Immunostaining for the scale IFE marker K31 was used to distinguish scale from interscale IFE. Using the free image processing software FIJI, a mask was created based on K31 immunostaining only considering depilated but otherwise intact scale:interscale units for the analysis. Immunostainings for Trp2 (MCs) and Langerin (LCs) were merged with the scale:interscale mask for further compartment-specific quantification of MC or LC numbers. For analysis of DETC distribution, a mask based on Langerin immunostaining was created and merged with γδTCR immunostaining. For analysis of MC, LC or DETC distribution in *Id2*Ctrl/KO, FVB/N, C57BL/6 WT and partially MC-free tail epidermis of C57BL/6 mice, percentages of the total MC, LC or DETC population per scale:interscale unit localizing to either scale or interscale IFE were calculated. For analysis of MC, LC or DETC distribution in control and K14ΔLef1 mice, the mask of the scale IFE (based on K31 immunostaining (MC,LC) or Langerin immunostaining (DETC)) was either decreased to 60% of its size to mark the center region of the scale IFE, or expanded to 110% to span the scale IFE and its periphery. Remaining K31-negative IFE was considered as the third zone (interscale). Percentages of the total MC, LC or DETC populations per scale:interscale unit localizing to one of the 3 defined regions was calculated per scale:interscale unit. Analysis of distribution and densities of MCs, LCs and DETCs comprised 2 to 15 scale:interscale units per animal.

### Analysis of apoptosis in tail epidermal whole mounts

For analysis of apoptotic MCs in K14ΔNLef1 mice and of apoptotic immune cells in *Lrig1KO* mice, partly depilated and acetone fixed tail epidermal whole mounts were blocked in PB buffer for 1 h at RT followed by incubation with primary antibodies diluted in PB buffer over-night at RT. The next day samples were washed three times for 10 min each with 0.05% Triton X- 100/PBS. Subsequently, samples were incubated with secondary antibodies (diluted in PB buffer) for 2 h at RT, followed by three washing steps (10 min each, RT) and mounting of samples in Mowiol, with the basal side facing upwards. Apoptotic cells in the remaining hair follicles served as internal positive control for cleaved Caspase3 immunostaining, as apoptosis is a natural process occurring during catagen.

### Quantitative analysis of MC, LC and DETC densities in tail epidermal whole-mounts

For analysis of MC, LC, or DETC densities in tail epidermis of *Id2*Ctrl/KO, FVB/N, C57BL/6 WT, and partially MC-free tail epidermis of C57BL/6 mice, respective cell numbers per scale or interscale compartment (determined by K31 (MC, LC) or Langerin immunostaining (DETC)) were counted and cell numbers per mm^2^ were calculated. For analysis of MC, LC or DETC densities in tail epidermis of C57BL/6 mice at P5, P10, and P21, respective cell numbers per complete scale:interscale unit were counted and cell numbers per mm^2^ were calculated. For analysis of MC, LC, or DETC densities in tail epidermis of control and K14ΔLef1 mice, respective cell numbers per compartment (scale center, scale periphery or interscale IFE, determined by K31 (MC, LC) or Langerin immunostaining (DETC) as previously described) were counted and cell numbers per mm^2^ were calculated.

### Quantitative analysis of MC and immune cell densities in ear epidermal whole-mounts

For quantification of MC, LC and DETC density in the ear epidermis, respective cell numbers were counted and numbers per mm^2^ were calculated. At least 0.825 mm^2^ per animal were analysed.

### Quantification of MC mean area and major axis length

A CellProfiler (Carpenter et al., 2006) pipeline was used to calculate MC mean area and major axis length. Briefly, Trp2 immunostaining was used for primary object identification. MC objects were assigned to tail IFE compartments (scale or interscale (C57BL/6), scale center or scale periphery or interscale IFE (K14ΔNLef1 mice)) and subsequent analysis was carried out automatically with supervision.

### Quantification of numbers of MC dendrites

MCs were assigned to tail IFE compartments (scale:interscale (C57BL/6), scale center or scale periphery or interscale IFE (K14ΔNLef1 mice)) and subsequently numbers of dendrites of MCs per compartment were determined.

### Antibodies

Details on the antibodies used in this study are listed in Table S1.

### Statistical analyses

Statistical analyses were performed using GraphPad Prism software (GraphPad, version 6.0). Significance was determined as indicated in the figure legends. N-values refer to biological replicates (=mice), are specified in the figure legends and correspond to the sample size used to derive statistics. All data sets were subjected to normality tests (D’Agostino–Pearson omnibus test, KS normality test, or Shapiro–Wilk normality test) when applicable. P-values are ranged as follows: *, p<0.05; **, p<0.01; ***, p<0.001; ****, p<0.0001 as detailed in the figure legends. For all experiments, measurements were taken from minimum three independent biological samples.

### Software

For data analysis, the following software has been used: GraphPad PRISM VI, Inkscape, ImageJ/Fiji (Rueden et al., 2017; Schindelin et al., 2012), Cell Profiler (Carpenter et al., 2006), ZEN 3.0 (blue edition; Carl Zeiss Microscopy GmbH, 2019).

## ACKNOWLEDGEMENTS

We thank the imaging facilities at CECAD and Saarland University (ALM-CF), and the imaging facility and UoC animal facilities for important services, and Florian Kuester and CMMC Imaging facility for assisting with mouse analysis. We are grateful to Wendy Havran (†) and Deborah Witherden for advice on DETC immunostaining. We acknowledge Kim B. Jensen (University of Copenhagen, Danstem, Denmark) for sharing Lrig1-CreERT2/EGFP mice. We thank all members of the Iden laboratory for stimulating discussions.

## DATA AVAILABILITY

Correspondence and requests for materials related to this study should be sent to. All data supporting the findings of this study are available within the paper and its supplemental information, or from the corresponding author on reasonable request.

## AUTHOR CONTRIBUTIONS

Conceptualization, methodology, validation: S.B., A.G., S.I.; investigation: S.B., A.K.B., S.C., S.I.; formal analysis: S.B., S.I.; resources: S.I.; visualization: S.B.; mouse models: S.I., C.N., K.S., M.Z.; writing (original draft): S.B., S.I.; writing (review, editing): S.I.; supervision and funding acquisition: S.I.. All authors provided input to the manuscript and data.

---

**Figure S1:**
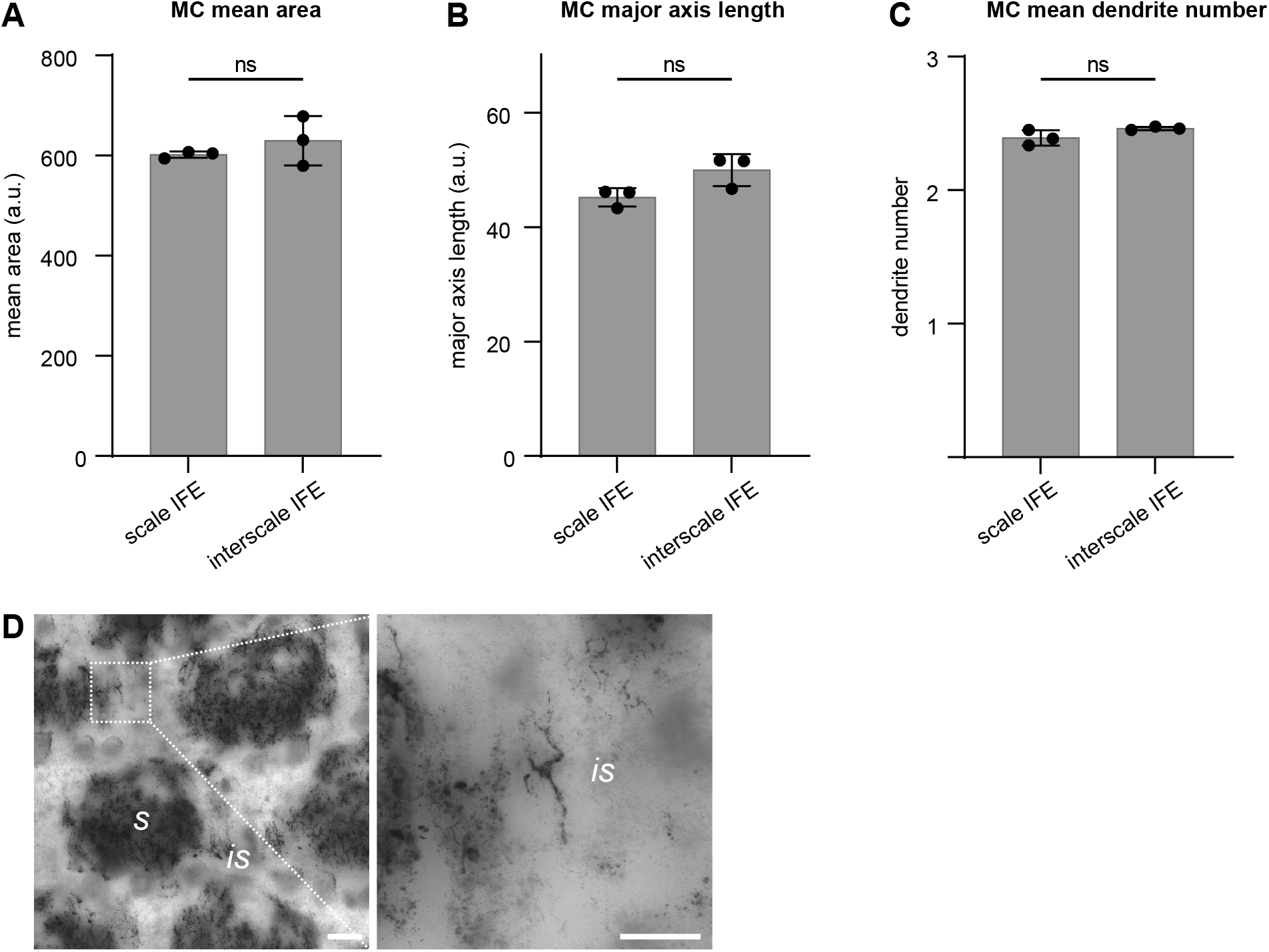
Interscale IFE contains a small MC population with morphological features similar to their scale-based counterparts and capable of melanin production. (A) Quantification of mean area per MC in 3-months old wild-type C57BL/6 scale and interscale tail IFE. n=3; ns: p=0.7000; mean±s.d.; Mann-Whitney test. (B) Quantification of major axis length per MC in 3-months old C57BL/6 scale and interscale tail IFE. n=3; ns: p=0.1000; mean±s.d.; Mann-Whitney test. (C) Quantification of dendrite numbers per MC in 3-months old C57BL/6 scale and interscale tail IFE. n=3; ns: p=0.2000; mean±s.d.; Mann-Whitney test. (D) Representative images of Fontana-Masson staining in tail epidermis whole-mounts of 3-months old C57BL/6 mice. Representative for n=3. Scale bars: 100 µm (right), 50 µm (left). s: scale, is: interscale.

**Figure S2:**
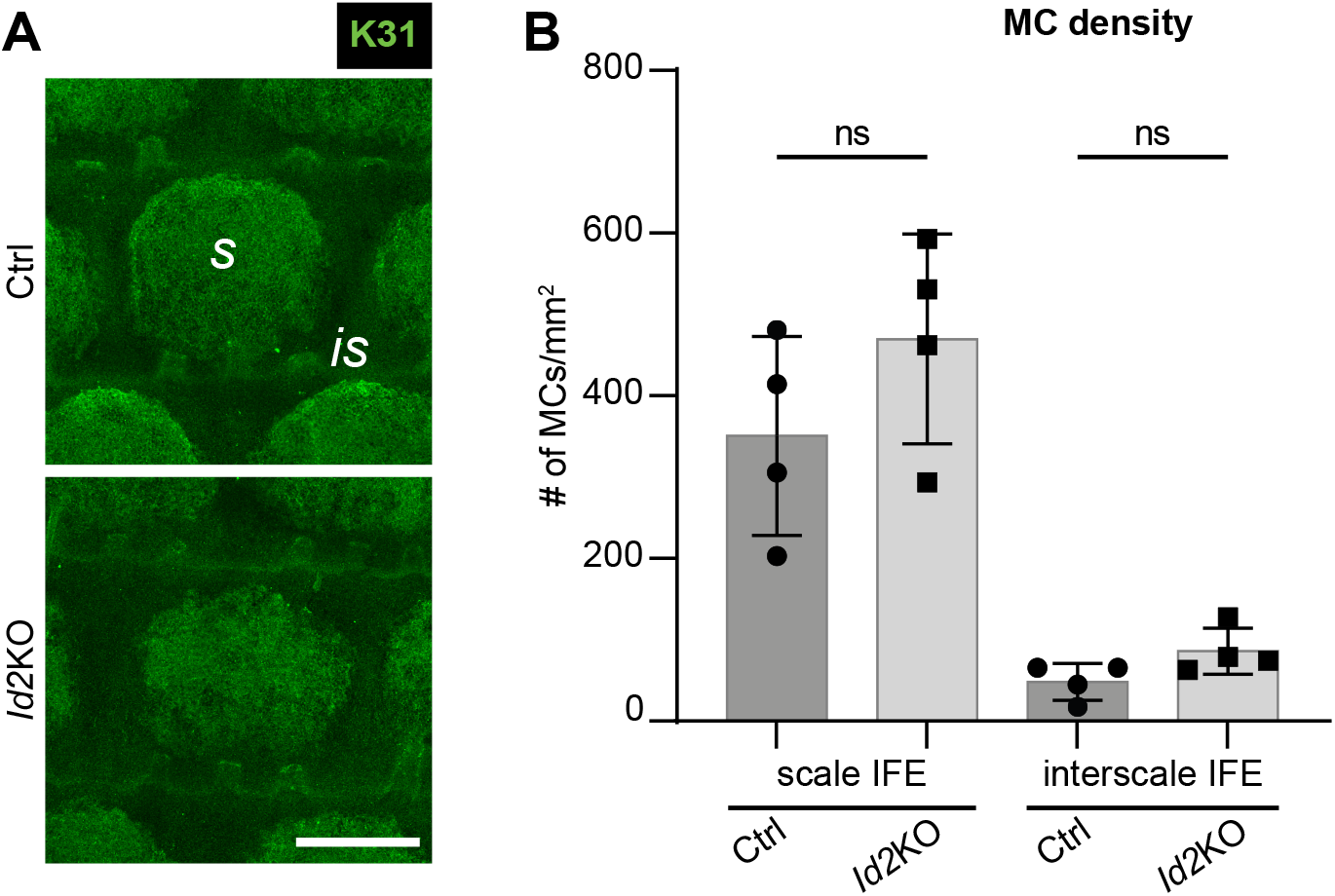
Epidermal scale differentiation and MC densities are unaffected by deletion of transcription factor Id2. (A) K31 immunostaining of tail epidermis whole-mounts of 3-months old control and *Id2*KO mice. Representative for n=4. Scale bar: 250 µm. (B) Quantification of 3A; MC cell numbers per scale:interscale unit (MCs/mm^2^) in tail epidermis from 3-months old control and *Id2*KO mice. n=4; ns: p=0.2962 (scale IFE, Ctrl vs. *Id2*KO); ns: p=0.9343 (interscale IFE, Ctrl vs. *Id2*KO); mean±s.d.; one-way ANOVA/Tukey’s multiple comparisons test. s: scale, is: interscale.

**Figure S3:**
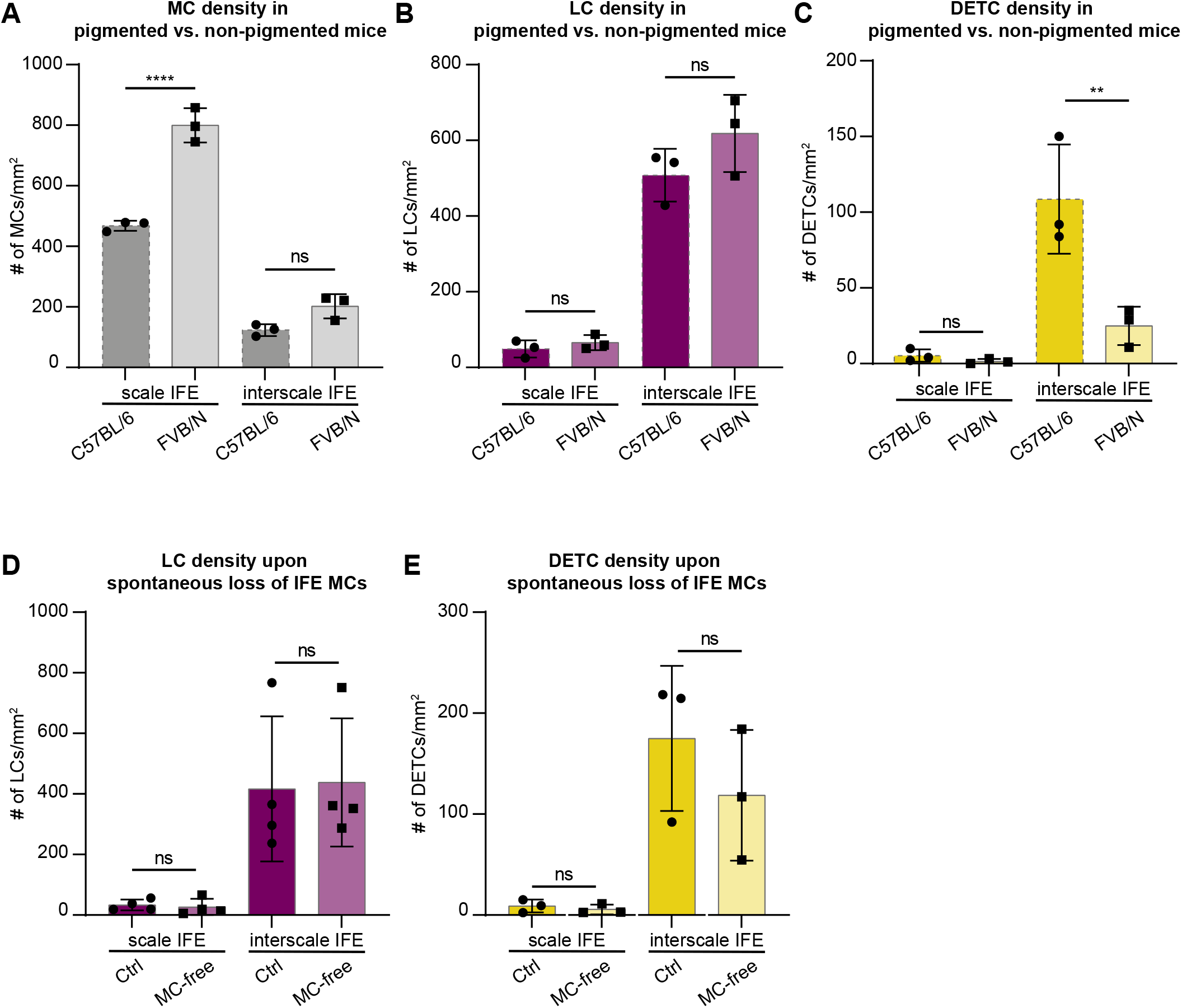
Epidermis-resident cell types in non-pigmented FVB/N mice and in regions of spontaneous MC loss. (A) Quantification of 4A; MC cell numbers per scale:interscale unit (MCs/mm^2^) in tail epidermis from 3-months old wild-type C57BL/6 and FVB/N mice. n=3; ****: p<0.0001 (scale IFE, C57BL/6 vs. FVB/N); ns: p=0.1173 (interscale IFE, C57BL/6 vs. FVB/N); mean±s.d.; one-way ANOVA/Tukey’s multiple comparisons test. Data for C57BL/6 as shown in Figure 1F. (B) Quantification of 4A; LC cell numbers per scale:interscale unit (LCs/mm^2^) in tail epidermis from 3-months old wild-type C57BL/6 and FVB/N mice. n=3; ns: p=0.9875 (scale IFE, C57BL/6 vs. FVB/N); ns: p=0.2228 (interscale IFE, C57BL/6 vs. FVB/N); mean±s.d.; one-way ANOVA/Tukey’s multiple comparisons test. Data for C57BL/6 as shown in Figure 1G. (C) Quantification of 4B; DETC cell numbers per scale:interscale unit (DETCs/mm^2^) in tail epidermis from 3-months old wild-type C57BL/6 and FVB/N mice. n=3; ns: p=0.9944 (scale IFE, C57BL/6 vs. FVB/N); **: p=0.0032 (interscale IFE, C57BL/6 vs. FVB/N); mean±s.d.; one-way ANOVA/Tukey’s multiple comparisons test. Data for C57BL/6 as shown in Figure 1H. (D) Quantification of 4G; LC cell numbers per scale:interscale unit (LCs/mm^2^) in tail epidermis from 3-months old wild-type C57BL/6 mice with spontaneous loss of MCs in posterior tail epidermis. n=4; ns: p>0.9999 (scale IFE, Ctrl vs. MC-free area); ns: p=0.9975 (interscale IFE, Ctrl vs. MC-free area); mean±s.d.; one-way ANOVA/Tukey’s multiple comparisons test. (E) Quantification of 4H; DETC cell numbers per scale:interscale unit (DETCs/mm^2^) in tail epidermis from 3-months old wild-type C57BL/6 mice with spontaneous loss of MCs in posterior tail epidermis. n=3; ns: p=0.9998 (scale IFE, Ctrl vs. MC-free area); ns: p=0.5202 (interscale IFE, Ctrl vs. MC-free area); mean±s.d.; one-way ANOVA/Tukey’s multiple comparisons test.

**Figure S4:**
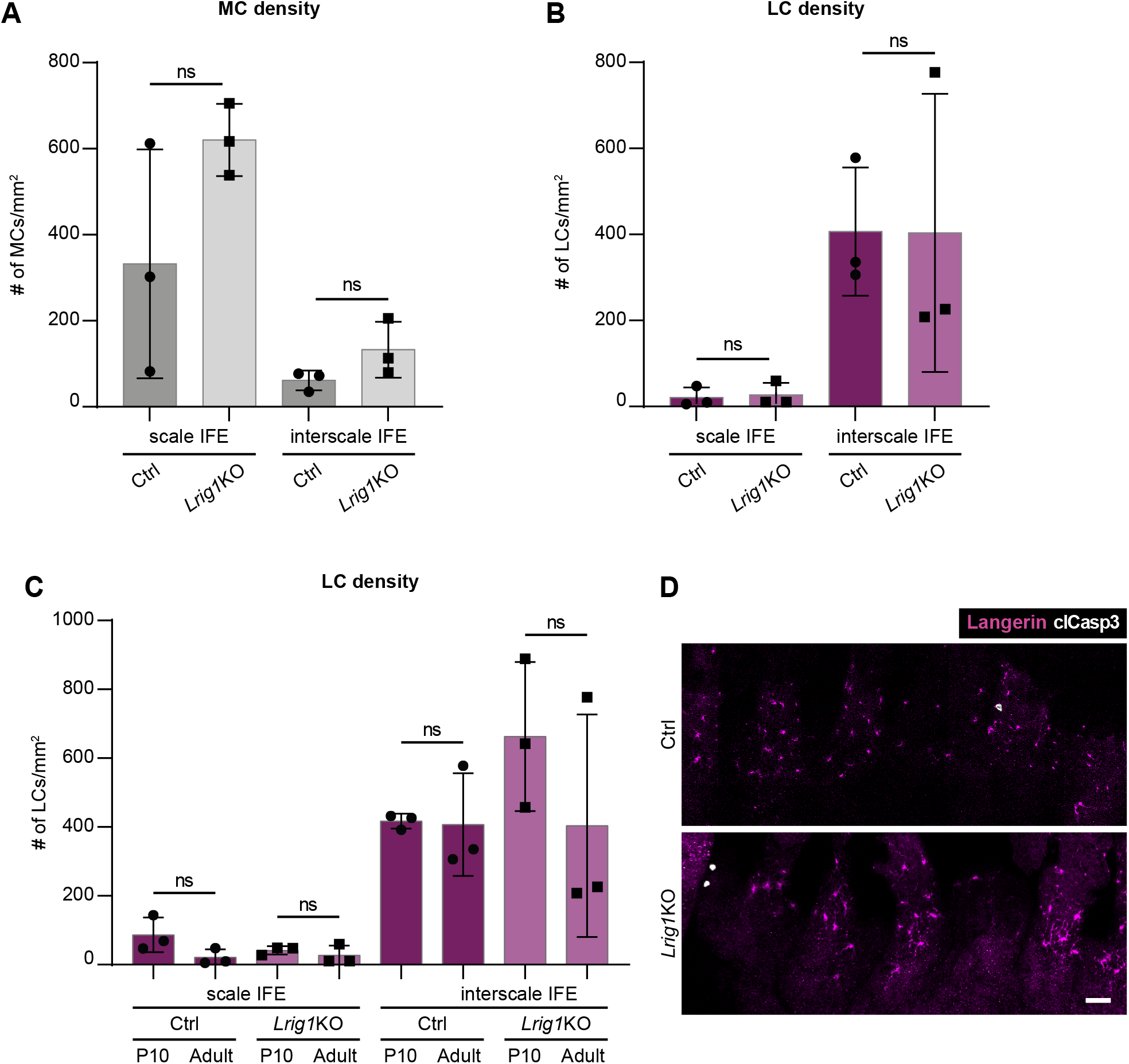
Analysis of resident cell types in *Lrig1*KO mice. (A) Quantification of 5A; MC cell numbers per scale:interscale unit (MCs/mm^2^) in tail epidermis from 3-months old control and *Lrig1*KO mice. n=3; ns: p=0.1435 (scale IFE, Ctrl vs. *Lrig1*KO); ns: p=0.9269 (interscale IFE, Ctrl vs. *Lrig1*KO); mean±s.d.; one-way ANOVA/Tukey’s multiple comparisons test. (B) Quantification of 5A; LC cell numbers per scale:interscale unit (LCs/mm^2^) in tail epidermis from 3-months old control and *Lrig1*KO mice. n=3; ns: p>0.9999 (scale IFE, Ctrl vs. *Lrig1*KO); ns: p>0.9999 (interscale IFE, Ctrl vs. *Lrig1*KO); mean±s.d.; one-way ANOVA/Tukey’s multiple comparisons test. (C) LC cell numbers per scale:interscale unit (LCs/mm^2^) in tail epidermis from P10 and 3-months old control and *Lrig1*KO mice (data set of 3-months old mice also shown in B, here compared to younger mice). n=3; ns: p=0.9992 (scale IFE, Ctrl P10 vs. Ctrl Adult); ns: p>0.9999 (scale IFE, *Lrig1*KO P10 vs. *Lrig1*KO Adult); ns: p>0.9999 (interscale IFE, Ctrl P10 vs. Ctrl Adult); ns: p=0.4413 (interscale IFE, *Lrig1*KO P10 vs. *Lrig1*KO Adult); mean±s.d.; one-way ANOVA/Tukey’s multiple comparisons test. (D) Langerin and cleaved Caspase3 immunostaining of tail epidermis whole-mounts from 3-months old control and *Lrig1*KO mice. Representative for n=3. Scale bar: 100 µm.

**Figure S5:**
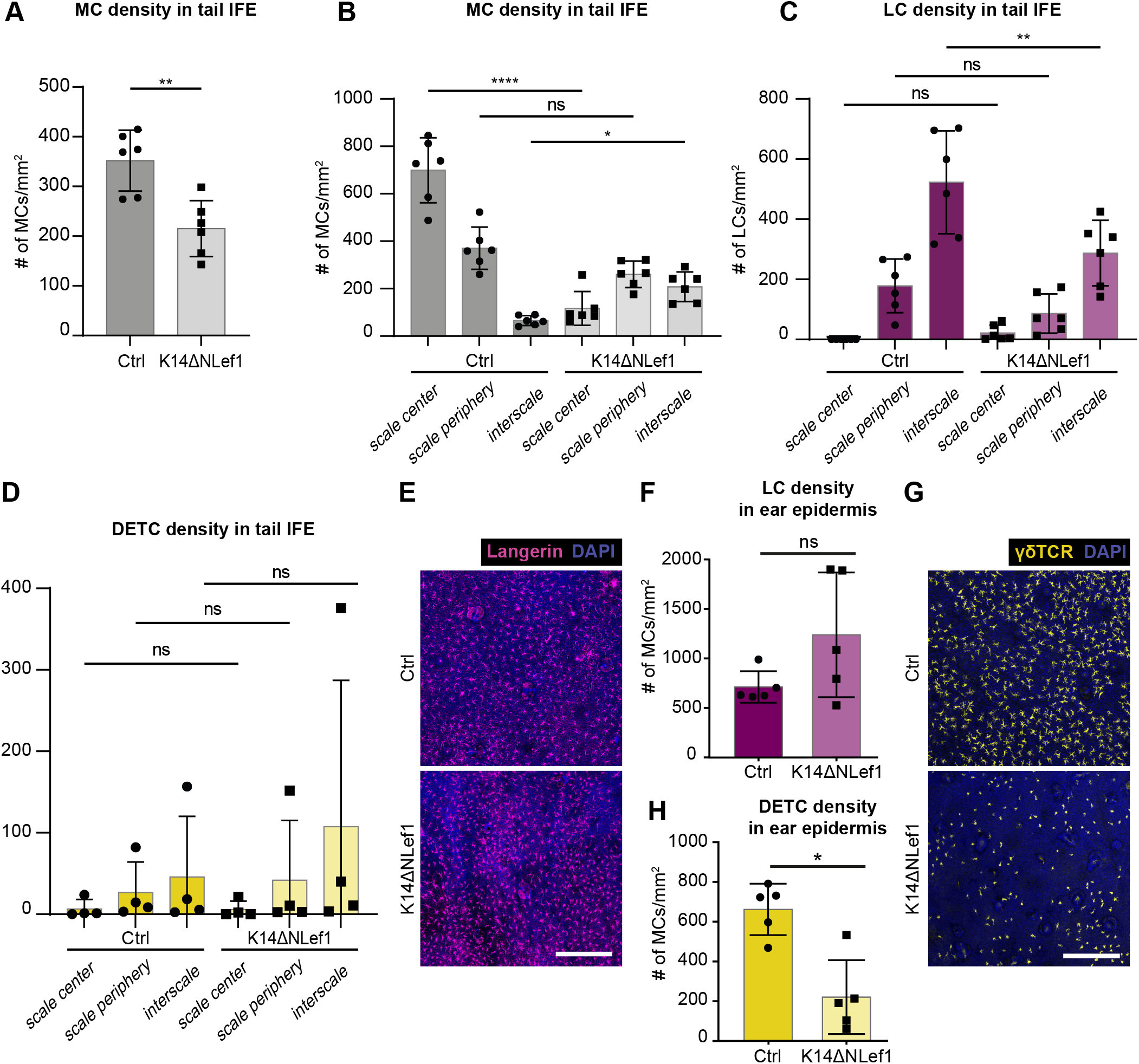
Analysis of resident cell types in K14ΔNLef1 tail and ear IFE. (A) Quantification of 6A; MC cell numbers (MCs/mm^2^) in tail epidermis from 3-months old control and K14ΔNLef1 mice. n=6; **: p=0.0087; mean±s.d.; Mann-Whitney test. (B) Quantification of 6A; MC cell numbers per scale:interscale unit (MCs/mm^2^) in tail epidermis from 3-months old control and K14ΔNLef1 mice. n=6; ****: p<0.0001 (scale center, Ctrl vs. K14ΔNLef1); ns: p=0.2070 (scale periphery, Ctrl vs. K14ΔNLef1); *: p=0.0497 (interscale IFE, Ctrl vs. K14ΔNLef1); mean±s.d.; one-way ANOVA/Tukey’s multiple comparisons test. (C) Quantification of 6A; LC cell numbers per scale:interscale unit (LCs/mm^2^) in tail epidermis from 3-months old control and K14ΔNLef1 mice. n=6; ns: p=0.9986 (scale center, Ctrl vs. K14ΔNLef1); ns: p=0.5590 (scale periphery, Ctrl vs. K14ΔNLef1); **: p=0.0021 (interscale IFE, Ctrl vs. K14ΔNLef1); mean±s.d.; one-way ANOVA/Tukey’s multiple comparisons test. (D) Quantification of 6B; DETC cell numbers per scale:interscale unit (DETCs/mm^2^) in tail epidermis from 3-months old control and K14ΔNLef1 mice. n=4; ns: p>0.9999 (scale center, Ctrl vs. K14ΔNLef1); ns: p>0.9999 (scale periphery, Ctrl vs. K14ΔNLef1); ns: p=0.8852 (interscale IFE, Ctrl vs. K14ΔNLef1); mean±s.d.; one-way ANOVA/Tukey’s multiple comparisons test. (E) Langerin immunostaining of ear epidermis whole-mounts of 3-months old control and K14ΔNLef1 mice. Nuclei were counterstained using DAPI. Scale bar: 250 µm. (F) Quantification of E; LC densities (number/mm^2^) in ear epidermis. n=5; ns: p=0.2222; mean±s.d.; Mann-Whitney test. (G) γδTCR immunostaining of ear epidermis whole-mounts of 3-months old control and K14ΔNLef1 mice. Nuclei were counterstained using DAPI. Scale bar: 250 µm. (H) Quantification of G; DETC densities (number/mm^2^) in ear epidermis. n=5; *: p=0.0159; mean±s.d.; Mann-Whitney test.

**Figure S6:**
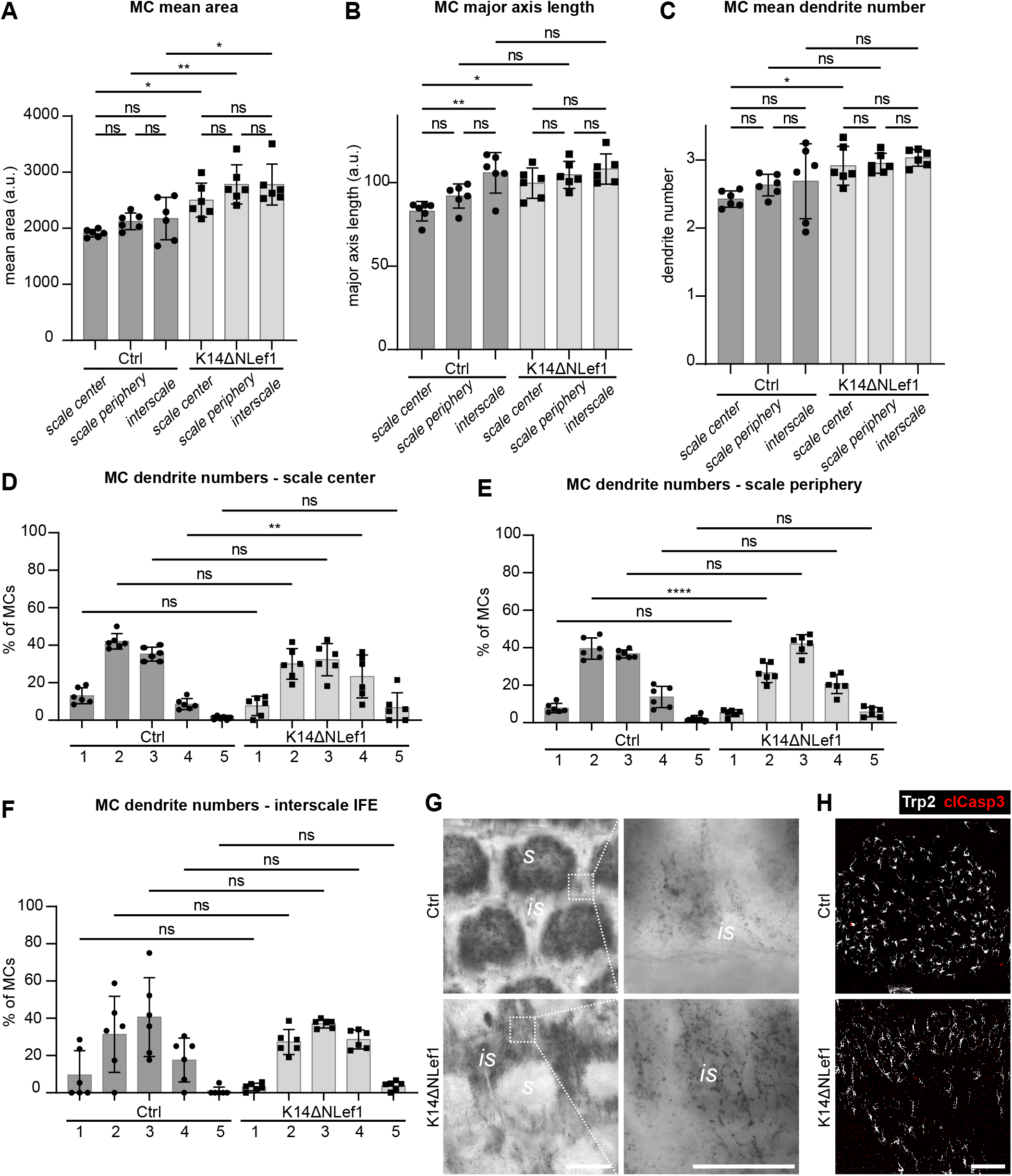
Analysis of MC properties in K14ΔNLef1 tail IFE. (A) Quantification of mean area per MC in tail IFE of 3-months old control and K14ΔNLef1 mice. n=6; ns: p=0.8155 (Ctrl, scale center vs. scale periphery); ns: p=0.6484 (Ctrl, scale center vs. interscale IFE); ns: p=0.9997 (Ctrl, scale periphery vs. interscale IFE); ns: p=0.5756 (K14ΔNLef1, scale center vs. scale periphery); ns: p=0.5801 (K14ΔNLef1, scale center vs. interscale IFE); ns: p>0.9999 (K14ΔNLef1, scale periphery vs. interscale IFE); *: p=0.0180 (scale center, Ctrl vs. K14ΔNLef1); **: p=0.0064 (scale periphery, Ctrl vs. K14ΔNLef1); *: p=0.0137 (intersale IFE, Ctrl vs. K14ΔNLef1); mean±s.d.; one-way ANOVA/Tukey’s multiple comparisons test. (B) Quantification of major axis length per MC in tail IFE of 3-months old control and K14ΔNLef1 mice. n=6; ns: p=0.4863 (Ctrl, scale center vs. scale periphery); **: p=0.0011 (Ctrl, scale center vs. interscale IFE); ns: p=0.0971 (Ctrl, scale periphery vs. interscale IFE); ns: p=0.9185 (K14ΔNLef1, scale center vs. scale periphery); ns: p=0.5608 (K14ΔNLef1, scale center vs. interscale IFE); ns: p=0.9825 (K14ΔNLef1, scale periphery vs. interscale IFE); *: p=0.0270 (scale center, Ctrl vs. K14ΔNLef1); ns: p=0.1156 (scale periphery, Ctrl vs. K14ΔNLef1); ns: p=0.9975 (intersale IFE, Ctrl vs. K14ΔNLef1); mean±s.d.; one-way ANOVA/Tukey’s multiple comparisons test. (C) Quantification of mean dendrite number per MC in tail IFE of 3-months old control and K14ΔNLef1 mice. n=6; ns: p=0.7946 (Ctrl, scale center vs. scale periphery); ns: p=0.5820 (Ctrl, scale center vs. interscale IFE); ns: p=0.9991 (Ctrl, scale periphery vs. interscale IFE); ns: p>0.9999 (K14ΔNLef1, scale center vs. scale periphery); ns: p=0.9765 (K14ΔNLef1, scale center vs. interscale IFE); ns: p=0.9957 (K14ΔNLef1, scale periphery vs. interscale IFE); *: p=0.0489 (scale center, Ctrl vs. K14ΔNLef1); ns: p=0.3645 (scale periphery, Ctrl vs. K14ΔNLef1); ns: p=0.2925 (intersale IFE, Ctrl vs. K14ΔNLef1); mean±s.d.; one-way ANOVA/Tukey’s multiple comparisons test. (D) Quantification of dendrite numbers (% of MCs) in tail IFE scale center compartment of 3-months old control and K14ΔNLef1 mice. n=6; ns: p=0.9083 (1 dendrite, Ctrl vs. K14ΔNLef1); ns: p=0.0635 (2 dendrites, Ctrl vs. K14ΔNLef1); ns: p=0.9983 (3 dendrites, Ctrl vs. K14ΔNLef1); **: p=0.0087 (4 dendrites, Ctrl vs. K14ΔNLef1); ns: p=0.8815 (5 dendrites, Ctrl vs. K14ΔNLef1); mean±s.d.; one-way ANOVA/Tukey’s multiple comparisons test. (E) Quantification of dendrite numbers (% of MCs) in tail IFE scale periphery compartment of 3-months old control and K14ΔNLef1 mice. n=6; ns: p=0.9790 (1 dendrite, Ctrl vs. K14ΔNLef1); ****: p<0.0001 (2 dendrites, Ctrl vs. K14ΔNLef1); ns: p=0-4488 (3 dendrites, Ctrl vs. K14ΔNLef1); ns: p=0.1162 (4 dendrites, Ctrl vs. K14ΔNLef1); ns: p=0.9074 (5 dendrites, Ctrl vs. K14ΔNLef1); mean±s.d.; one-way ANOVA/Tukey’s multiple comparisons test. (F) Quantification of dendrite numbers (% of MCs) in tail IFE interscale compartment of 3-months old control and K14ΔNLef1 mice. n=6; ns: p=0.9910 (1 dendrite, Ctrl vs. K14ΔNLef1); ns: p=0.9998 (2 dendrites, Ctrl vs. K14ΔNLef1) ns: p>0.9999 (3 dendrites, Ctrl vs. K14ΔNLef1); ns: p=0.7885 (4 dendrites, Ctrl vs. K14ΔNLef1); ns: p>0.9999 (5 dendrites, Ctrl vs. K14ΔNLef1 mean±s.d.; one-way ANOVA/Tukey’s multiple comparisons test. (G) Representative micrographs of Fontana-Masson staining in tail epidermis whole-mounts of 3-months old Ctrl and K14ΔNLef1 mice. Representative for n=3. Scale bars: 300 µm (left), 100 µm (right). (H) Langerin and cleaved Caspase3 immunostaining of tail epidermis whole-mounts from 3-months old control and K14ΔNLef1 mice. Representative for n=3. Scale bar: 100 µm. s: scale, is: interscale.

**Supplementary Table 1:** Antibodies and imaging reagents used in this study.

| **Primary antibodies** |  |  |  |  |
| --- | --- | --- | --- | --- |
| **antibody (supplier)** | **catalog nr.** | **clone/Ref.** | **species** | **dilution** |
| K31 (Progen) | GP-HHA1 | polyclonal | guinea pig | IF 1:100 |
| Trp2 (Santa Cruz Biotechnology) | sc-10451 | polyclonal | goat | IF 1:100 |
| Langerin (Invitrogen) | 14-2073-80 | eBioRMUL.2 | rat | IF 1:100 |
| MHC II (M. Pasparakis lab, CECAD, Cologne) | - | - | rat | IF 1:100 |
| CD45 (Invitrogen) | 14-0451-82 | 30-F11 | rat | IF 1:100 |
| Cleaved caspase 3 (R&D Systems) | AF835 | - | rabbit | IF 1:100 |
| **Secondary antibodies** |  |  |  |  |
| Goat anti-guinea pig AF488 (Invitrogen) | A11073 | polyclonal | goat | IF 1:500 |
| Donkey anti-rat AF594 (Dianova) | 712-585-153 | polyclonal | donkey | IF 1:500 |
| Donkey anti-goat AF647 (Invitrogen) | A21447 | polyclonal | donkey | IF 1:500 |
| Goat anti-rat AF647 (Invitrogen) | A21247 | polyclonal | goat | IF 1:500 |
| Goat anti-rabbit AF568 (Invitrogen) | A11036 | polyclonal | goat | IF 1:500 |
| Donkey anti-goat AF594 (Invitrogen) | A11058 | polyclonal | donkey | IF 1:500 |
| Donkey anti-rabbit AF488 (Invitrogen) | A21206 | polyclonal | donkey | IF 1:500 |
| Goat anti-guinea pig AF568 (Invitrogen) | A11075 | polyclonal | goat | IF 1:500 |
| **Other reagents** |  |  |  |  |
| DAPI (Carl Roth) | - | - | - | IF 1:500 |
| Fontana-Masson staining kit (Sigma-Aldrich) | HT200 | - | - | - |
| **Directly-conjugated antibodies** |  |  |  |  |
| FITC anti-mouse TCR $\gamma/\delta$ (BioLegend) | 118105 | GL3 | Armenian hamster | IF 1:300 |

